# Microbial communities increase host plant leaf growth in a pitcher plant experimental system

**DOI:** 10.1101/2024.01.30.578016

**Authors:** Jessica R. Bernardin, Erica B. Young, Sarah M. Gray, Leonora S. Bittleston

## Abstract

Across diverse ecosystems, bacteria and their host organisms engage in complex relationships having negative, neutral, or positive interactions. However, the specific effects of leaf-associated bacterial community functions on plant growth are poorly understood. To address this gap, we explored the relationships between bacterial community function and host plant growth in the purple pitcher plant (*Sarracenia purpurea*). The main aim of our research was to investigate how different bacterial community functions affect the growth and nutrient content in the plant. Previous research had suggested that microbial communities may aid in prey decomposition and subsequent nutrient acquisition in carnivorous plants, including *S. purpurea*. However, the specific functional roles of these bacterial communities in plant growth and nutrient uptake are not well known. In this study, sterile, freshly opened leaves (pitchers) were inoculated with three functionally distinct, pre-assembled bacterial communities and effects examined over 8 weeks. Bacterial community composition and function were measured using physiological assays, metagenomics, and metatranscriptomics. Distinct bacterial functions affected plant traits; a bacterial community enriched in decomposition and secondary metabolite production traits was associated with larger leaves with almost double the biomass of control pitchers. Physiological differences in bacterial communities were supported by metatranscriptomic analysis; for example, the bacterial community with the highest chitinase activity had greater expression of transcripts associated with chitinase enzymes. The relationship between bacterial community function and plant growth observed here indicates potential mechanisms for host-associated bacterial functions to support plant health and growth.

**Importance:** This study addresses a gap in understanding the relationships between bacterial community function and plant growth. We inoculated sterile, freshly opened pitcher plant leaves with three functionally distinct bacterial communities to uncover potential mechanisms through which bacterial functions support plant health and growth. Our findings demonstrate that distinct bacterial functions significantly influence plant traits, with some bacterial communities supporting more growth than in control pitchers. These results highlight the ecological roles of microbial communities in plants and thus ecosystems, and suggest potential pathways in which microbes support host plant health. This research provides valuable insights into plant-microbe interactions and effects of diverse microbial community functions.

## Introduction

Microbiomes have strong effects on the health and fitness of their hosts (1, 2). Host plant growth and physiology are influenced by the presence of leaf-associated (phyllosphere) microbes, with microbial taxa acting as mutualists, commensals, and antagonistic pathogens (3–6). Phyllosphere microbes live in complex microecosystems (7, 8) performing functions that contribute to plant nutrient acquisition, phytohormone production, and stress tolerance (3).

For example, the addition of synthetic phyllosphere communities was shown to increase fruit production in greenhouse-grown tomatoes (6) and increase leaf sugar content and metabolic diversity in field-grown *Agave tequilana* (9). Plants reap many benefits from hosting microbial communities (3, 9). Nutrient cycling interactions between microbes and plants can occur directly, e.g., via bacterial nitrogen fixation, and indirectly, e.g., via increased bacterial growth that influences host plant metabolism (3, 10). Many bacteria associated with plant-microbe interactions also have pathways to produce phytohormones, like auxin or cytokinins, which support plant cell metabolism and growth (3, 11–15). Plant growth-promoting bacteria can also mediate host stress responses, including increasing drought tolerance, thermotolerance, and UV protection (3, 16, 17). Despite these demonstrated benefits to host plants, we still lack research directly connecting bacterial community function with host physiological functions and benefits. We define bacterial community function as the collective metabolic processes of all the individual bacteria within a community. We addressed this knowledge gap by examining both bacterial and plant physiology, using metabarcoding, metagenomics, and metatranscriptomics of bacterial communities and measuring plant responses.

The carnivorous pitcher plant, *Sarracenia purpurea* L., uses water-filled, cup-shaped leaves to trap and digest insects to supplement acquisition of nutrients from low nutrient soils (18, 19). These leaves or ‘pitchers’ host an aquatic micro-ecosystem composed of microbes, fungi, protists, and invertebrates (8, 20, 21). The microbes are thought to provide critical ecosystem functions to the host plants by mediating insect prey digestion through the production of extracellular hydrolytic enzymes. It is likely that the bacterial production of protease and chitinase enzymes that digest protein and chitin-rich insect prey are especially important for this carnivorous plant species (22–24). *Sarracenia purpurea* pitcher plants do not produce chitinase enzymes (22). Thus, they may rely heavily on the bacterial extracellular enzymes to catalyze decomposition of insect prey and support the mineralization of carbon, nitrogen, and other nutrients for the host plant and the whole micro-ecosystem. This is an ideal model system to examine bacterial functions in controlled manipulative experiments, serving as an analog for other systems where experimentation is more challenging. Although previous research has hypothesized that pitcher plant leaf-associated bacterial communities benefit the host plant (18, 25, 26), no studies have specifically measured a direct benefit to the plant.

To connect the effects of bacterial function to plant growth, we designed a manipulative experiment using pitcher plant leaves and bacterial communities with different physiological functional profiles, measured as different levels of activity across different physiological functions (Fig. S1). These community physiological functions, including hydrolytic enzyme activity, organic substrate use, and respiration rates, were characterized for the three distinct experimental bacterial communities prior to inoculation, and the effects of these functionally distinct bacterial communities on host plants, were examined over eight weeks (Fig. 1).

**Figure 1.**
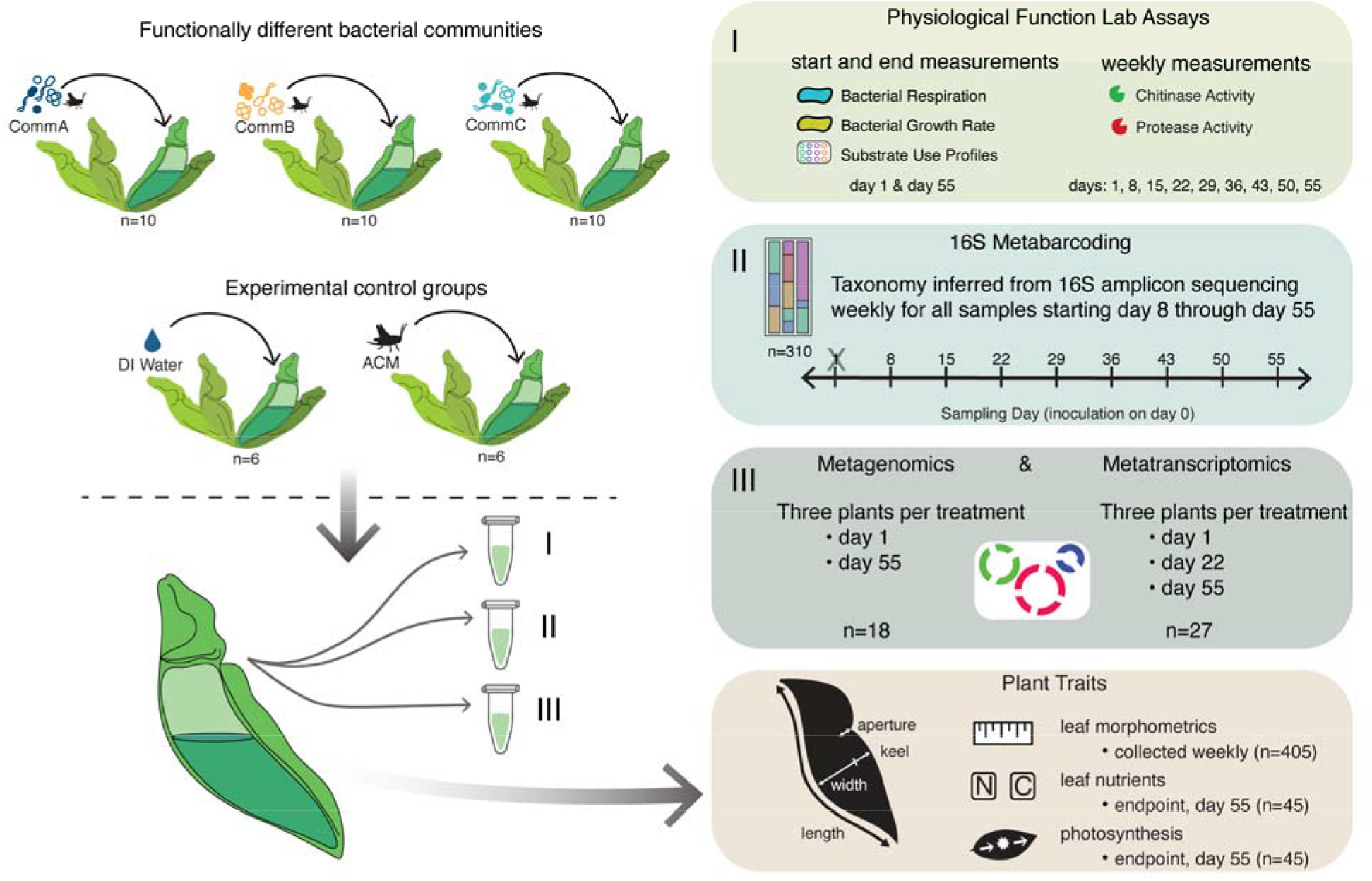
Experimental design. Three functionally distinct bacterial communities (CommA, CommB, and CommC) growing in sterile acidified cricket media (ACM) were inoculated into sterile pitchers and plants incubated in growth chambers for eight weeks. Sterile deionized (DI) water or ACM were added to additional pitchers as experimental controls. Pitcher fluid samples were collected weekly to characterize bacterial community composition and function: chitinase and protease activity, community respiration, growth rate, organic substrate use profiles, 16S amplicon sequencing, metagenomics, and metatranscriptomics. Plant functional traits were also assessed, including morphometric measurements of pitchers, as well as end-of-experiment measures of leaf biomass, leaf nutrient content, and photosynthetic rate.

We asked four specific questions:

1. Do bacterial communities with different functional profiles have differential effects on plant traits, such as leaf growth and nutrients?
2. Are there taxonomic characteristics of these bacterial communities that can explain the observed functional differences?
3. Can bacterial functions directly measured with physiological assays be linked with differential gene expression (metatranscriptomics) to better understand how bacterial community function impacts their host?
4. Are there specific functions, e.g., hydrolytic enzyme activity or carbon substrate use, that drive these effects on host plants?

We hypothesized that communities with specific physiological functions (e.g., high chitinase activity) would differentially express genes related to degradation, helping to release and make available more nutrients to enhance host plant growth.

## Results

### Starting communities were functionally different

The three communities (CommA, CommB, and CommC) inoculated into pitchers were functionally different in their chitinase activity, protease activity, bacterial respiration, and/or EcoPlate organic carbon substrate use profiles (Fig. 2, Fig. S2). Prior to inoculation, CommB had the highest chitinase activity and lowest protease activity (Fig. 2A, B, statistical differences shown in Fig. S2), and bacterial respiration was lower in CommB than the other two communities (Fig. S2). Using a permutational multivariate analysis of variance (PERMANOVA), we observed strong differences in carbon substrate use profiles among the three bacterial communities, both in the community cultures prior to inoculation (day 0) and when resampled from the pitchers 24 hours after inoculation (day 1) (Fig. 2C, Fig. S3, Table S1). CommB was the most distinct, while CommA and CommC exhibited similar levels of hydrolytic enzyme activity, all three communities had distinct carbon substrate use profiles. We found significant differences in community composition between the treatments on day 1 based on metagenomic sequencing (Fig. 2D, PERMANOVA; R^2^=0.78, F =10.382, P=0.005). There were no measured differences in bacterial community growth rates prior to inoculation or at any point during the experiment (Fig. S2).

**Figure 2.**
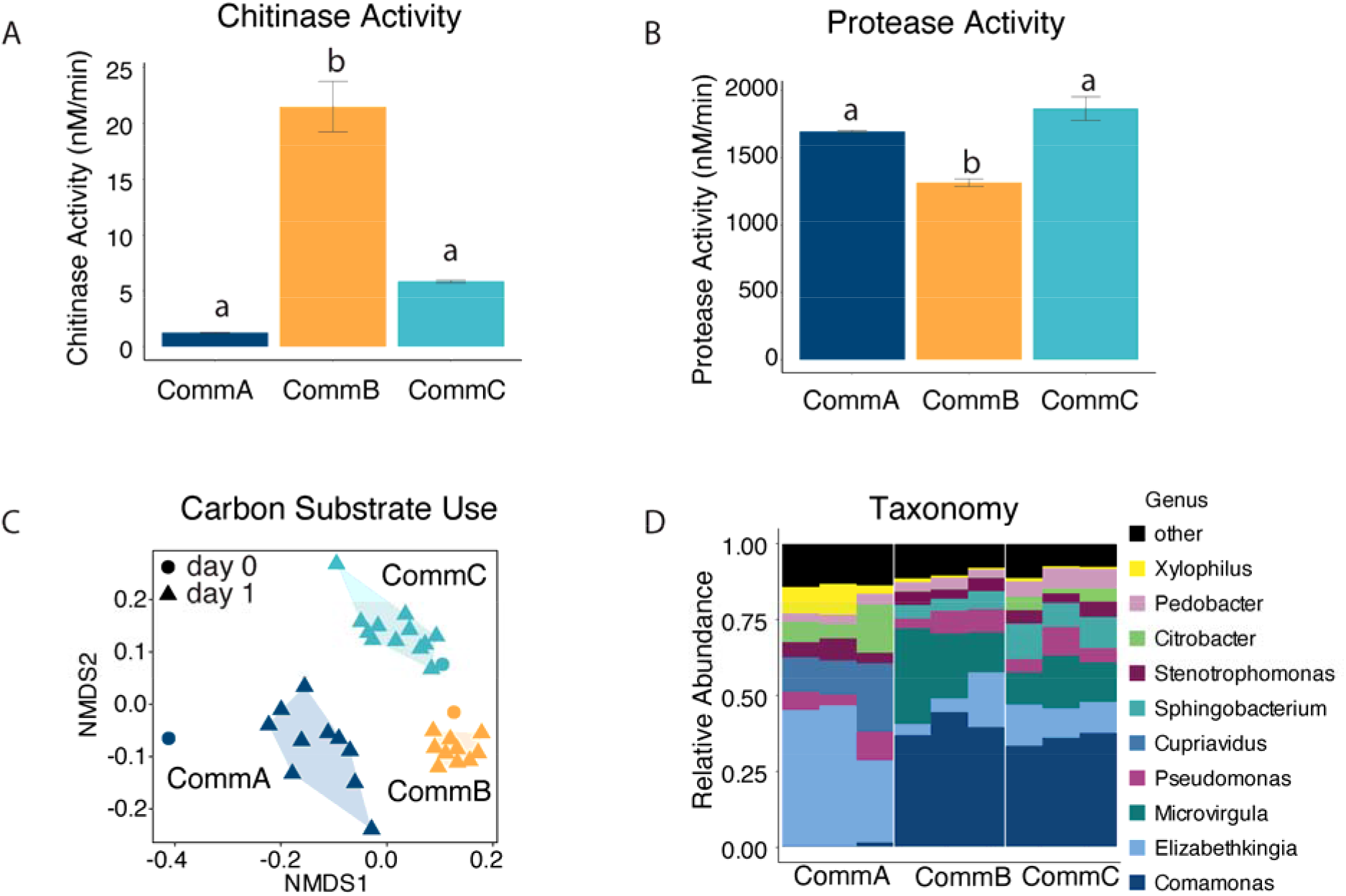
The three bacterial communities had different physiological functions measured before pitcher inoculation. (**A**) CommB had higher chitinase activity and (**B**) lower protease activity than CommA and CommC, there were no differences in bacterial growth rates between the starting treatments and CommB had the lowest bacterial respiration rates (Fig. S2). (**C**) The three communities also showed different EcoPlate carbon substrate use profiles, using Bray-Curtis dissimilarity and visualized with a NMDS (k=2, stress=0.138) at day 0 prior to inoculation and on day 1, 24 hours after inoculation. (**D**) These three communities also had distinct bacterial compositions (top 10 genera shown) at day one, based on taxonomy from shotgun metagenomic sequencing.

### Bacterial function affects plant traits

To investigate the influence of functionally diverse bacterial communities on plant traits (Question 1), we evaluated how experimental treatments (CommA, CommB, and CommC) and control treatments (water and ACM) affected pitcher biomass and nitrogen content after the plants hosted the treatments for an 8-week period. We compared our treatment communities to two control groups to help parse out differences between the effect of hosting the bacterial community vs. the media they are grown in, since pitcher plants may be able to directly absorb amino acids from degraded prey items (27). Pitchers hosting CommB had 1.7 times higher biomass (mean biomass = 0.47 g) than pitchers with cricket media only (mean biomass = 0.28 g) and 2 times higher biomass than the water-only plants (mean biomass = 0.23 g, Fig. 3A, 3B). We observed that plants that hosted CommB had higher biomass (Fig. 3A, 3B), longer pitcher length, and higher carbon content (Fig. S4) than not only the water and media controls, but also the other experimental treatments, CommA and CommC. While pitchers size varied and the sample size was relatively low due to the slow growth of this perennial plant (n=46), the probability of observing a higher pitcher biomass in bacterial community treatments compared to the cricket media (ACM) control was estimated at 81.0% for CommA, 99.7% for CommB, and 84.3% for CommC (based on posterior probabilities from GLMM, Fig. 3B). Although the 95% credible intervals (CIs) contained zero for CommC, we also saw a trend of a positive effect on total plant growth (measured as the net production of leaves). Plants in the CommC group had an estimated 1.4 greater net (gross new leaves - gross leaf loss) new pitcher production compared to the ACM control over eight weeks (Fig. S4).

**Figure 3.**
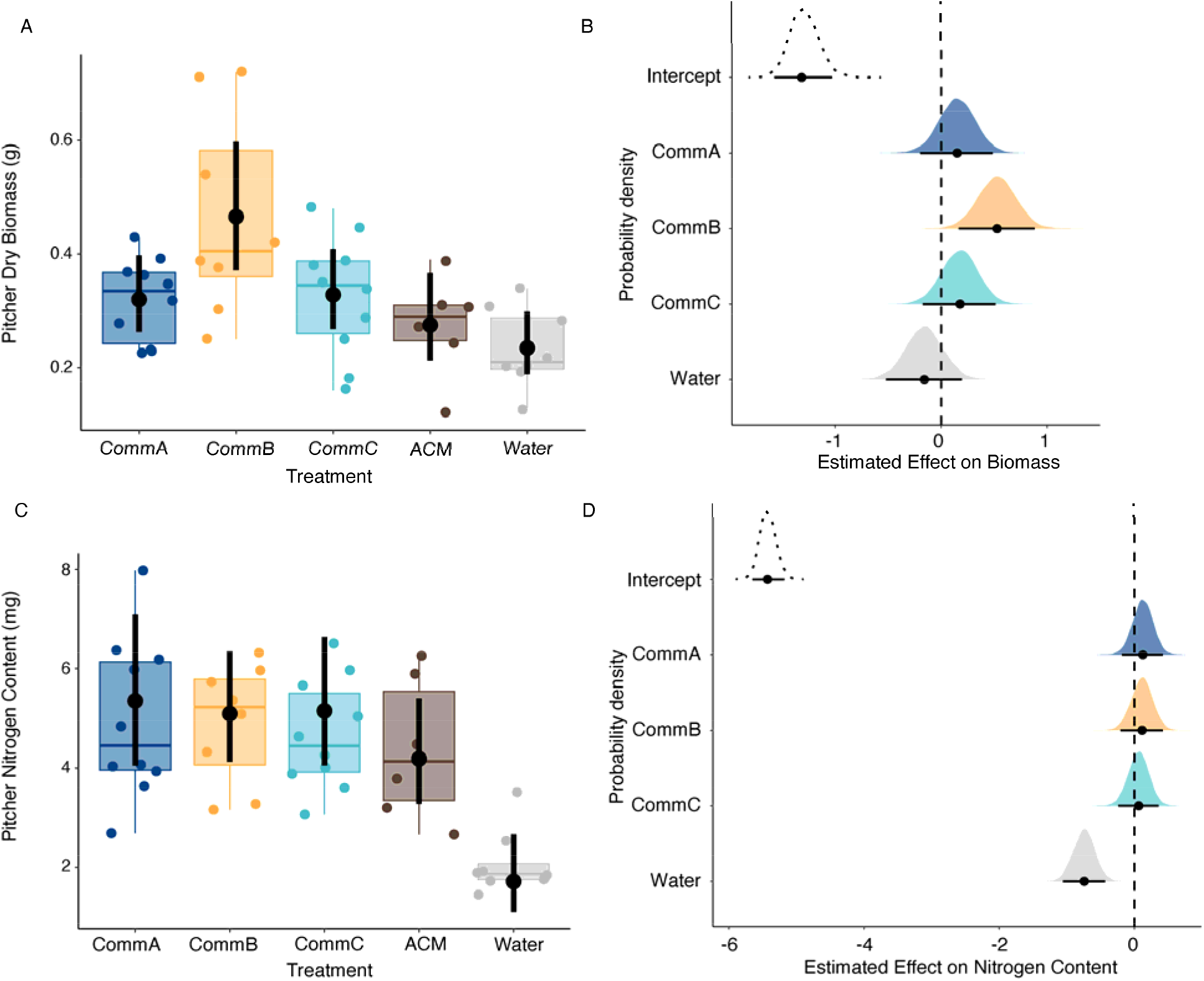
Bacterial community treatment affects plant traits. (**A**) Bayesian GLMM marginal effects of treatment on target pitcher biomass are represented by the black points, which are the median estimates, and the vertical lines represent the 95% credible intervals (CIs). The pitcher biomass data used to calculate estimated effects are represented by the box plot and colored points behind the marginal effect estimates. (**B**) Predicted effects of functionally distinct community treatments on pitcher biomass. Black points represent median estimates, lines represent the 95% CIs, and color-coded (by treatment) density plots indicate the full posterior. The intercept (dashed posterior) is set as the ACM baseline. (**C**) Bayesian GLMM marginal effects of treatment on pitcher nitrogen content, and (**D**) predicted effects of bacterial community treatments on pitcher nitrogen content.

Bacterial communities and the ACM control also affected nitrogen levels in pitcher plants (Fig. 3C and 3D). Comparing total pitcher nitrogen content between the treatment groups and the two controls (ACM, water), there was a 74% probability that bacterial treatments had higher nitrogen content than the ACM treatment, although the 95% credible intervals included zero. Water had a strong negative effect on pitcher nitrogen content, with plants containing only water having an estimated 1.7 mg of nitrogen, those with ACM having 4.2 mg, and those in the three bacterial treatments averaging 5.2 mg (Fig. 3C and 3D). We found no impact of the bacterial community treatment groups on other plant traits, including photosynthetic rates and photosynthetic quantum yield (F_v_/F_m_) (Fig. S4).

### Compositional differences between the three communities — 16S metabarcoding results

After quality control and rarefaction, we recovered 917 amplicon sequence variants (ASVs) in a community matrix spanning 310 samples (6-10 pitchers per treatment across 8 weekly time points). These ASVs represented 26 bacterial phyla, 192 families, and 286 genera. Among the 20 most abundant genera averaged across pitchers within a treatment at a single time point, *Microvirgula* (14.2%), *Stenotrophomonas* (12.4%), and *Pedobacter* (11.3%) accounted for almost 25% of reads (Fig. 4A).

**Figure 4.**
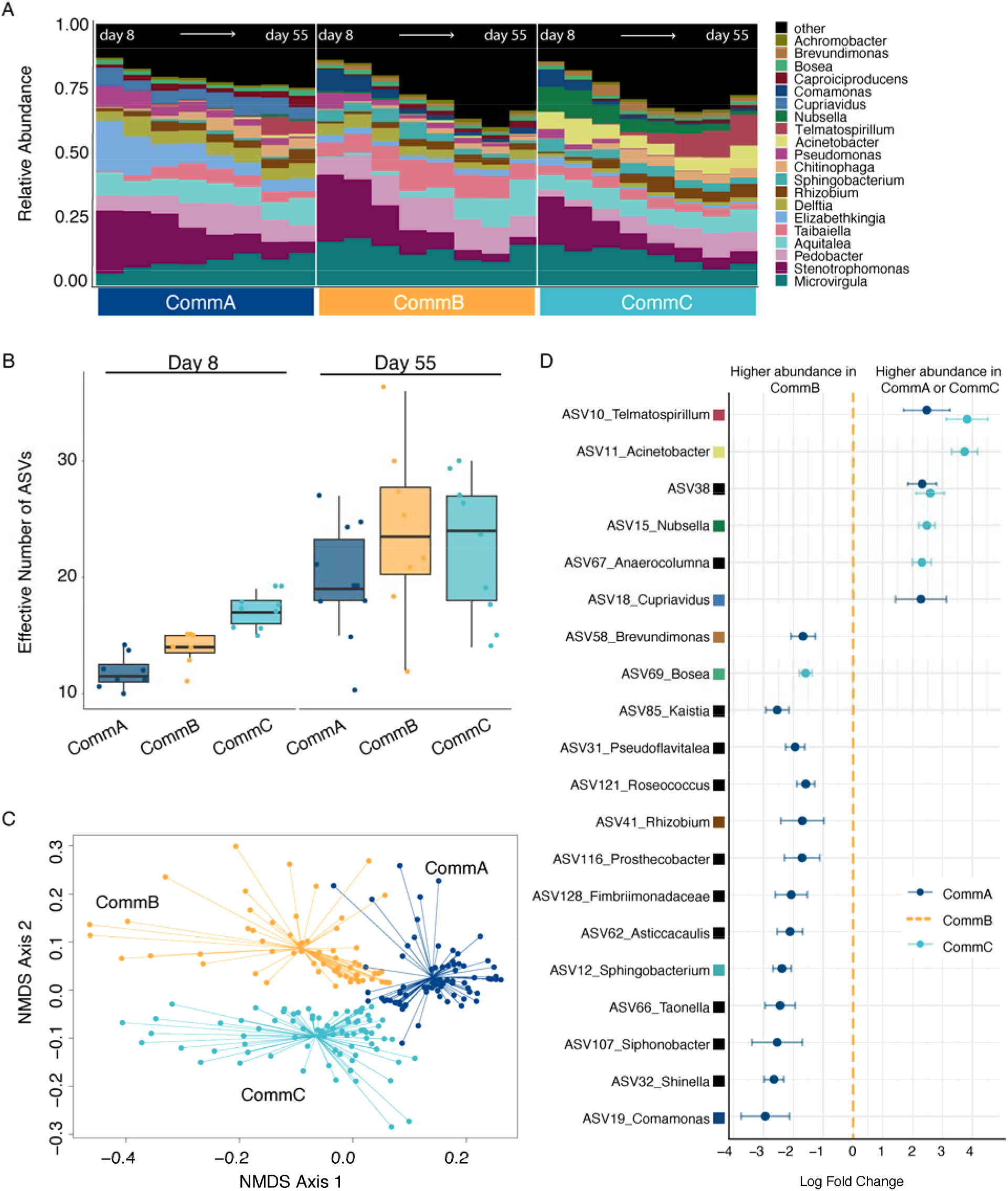
Bacterial community compositional differences between the three bacterial community treatments, based on 16S amplicon sequencing. (**A**) Relative abundance of the 20 most abundant genera from day 8 to 55 (days: 8, 15, 22, 29, 36, 43, 50, 55), with reads averaged for 10 different pitchers at the same time point. (**B**) Effective number of ASVs (Hill number 1) for each treatment at day 8 and day 55; box plots show the interquartile range, the horizontal lines are the median values and the whiskers extend 1.5 times the interquartile range, and each point represents a pitcher fluid sample. (**C**) Composition-based NMDS calculated using unweighted UniFrac for all samples within each of the three treatments from day 8 to 55, stress=0.17, k=2. (**D**) Twenty differentially abundant ASVs, predicted from ANCOM- BC2 (subset of ASVs with a log fold change greater than +/- 1.5) with CommB as the reference (orange dashed line). Points are colored by treatment and represent the log fold change of the model effects, with whiskers representing the standard error around each estimate. ASVs are labeled with assigned genus when available, and each colored square matches the color key in (**A**).

### Community Diversity and Differential Abundance

Addressing Question 2 with regard to taxonomic characteristics of the bacterial communities that could explain functional differences, we first observed differences in alpha diversity between the three treatments after one week *in planta* (Fig. 4B, Fig. S5). Between days 8 and 55, the effective number of ASVs increased (with a mean gain of 7.5 ASVs per sample) and by day 55, the differences in ASV numbers between community treatments had disappeared (Fig. 4B, Fig. S5). Beta diversity was different between the three community treatments (PERMANOVA; R^2^=0.282, F =42.04, p=0.001) using unweighted UniFrac distances, shown as clear clustering of treatments in NMDS visualization (Fig. 4C) and significant pairwise differences over time (Table S2). Mantel tests (Spearman’s rank correlation rho) showed significant correlations between the unweighted UniFrac distances of 16S community diversity with chitinase activity (r=0.227, p=0.001) and with final pitcher length (r=0.117, p=0.001), but no significant relationship with protease activity (r=0.004, p=0.44) or pitcher dry biomass (r=0.166, p=0.066).

Analysis of compositions of microbiomes with bias correction (ANCOM-BC) (28) identified 38 differentially abundant taxa between treatments (Table S3) with log fold changes (LFC) greater than 1.5 between at least one pair of treatments (CommB vs. CommA, CommC vs. CommA, or CommB vs. CommC). These taxa were plotted along with error estimates (Fig. 4D). In our model, CommB is set as the reference, so negative log fold change values indicate increased abundance in CommB (Fig. 4D, left side). CommB had a higher abundance of 13 taxa, with 12 of these taxa showing higher abundance in CommB compared to CommA, and 1 taxon showing higher abundance in CommB compared to CommC. Conversely, CommB had significantly lower abundance in 6 other taxa (Fig. 4D).

### Linking bacterial physiology with agnostic estimates of functional expression

To address Question 3, we identified 218,817 transcripts from 27 metatranscriptomic samples, including 3 pitchers each for 3 bacterial treatments at 3 different time points (Figure 1). This analysis links bacterial functions directly with differential gene expression (metatranscriptomics). After filtering out transcripts lacking KEGG orthologs, 162,624 total transcripts remained. We visualized the differences in metatranscriptomic profiles between the three bacterial treatments over time using an NMDS based on Bray-Curtis dissimilarities for all transcripts (Fig. 5A). We found clear differences between treatments (PERMANOVA; R^2^=0.20 F _22_=4.46, p=0.001), despite the many shared transcripts, and clear patterns through time (PERMANOVA; R^2^=0.27, F =6.00, p=0.001). The transcript by sample matrix of ∼162,000 genes was normalized and put into our differential expression (DE) model, which identified 24,357 unique significantly differentially expressed genes between our three bacterial communities.

**Figure 5.**
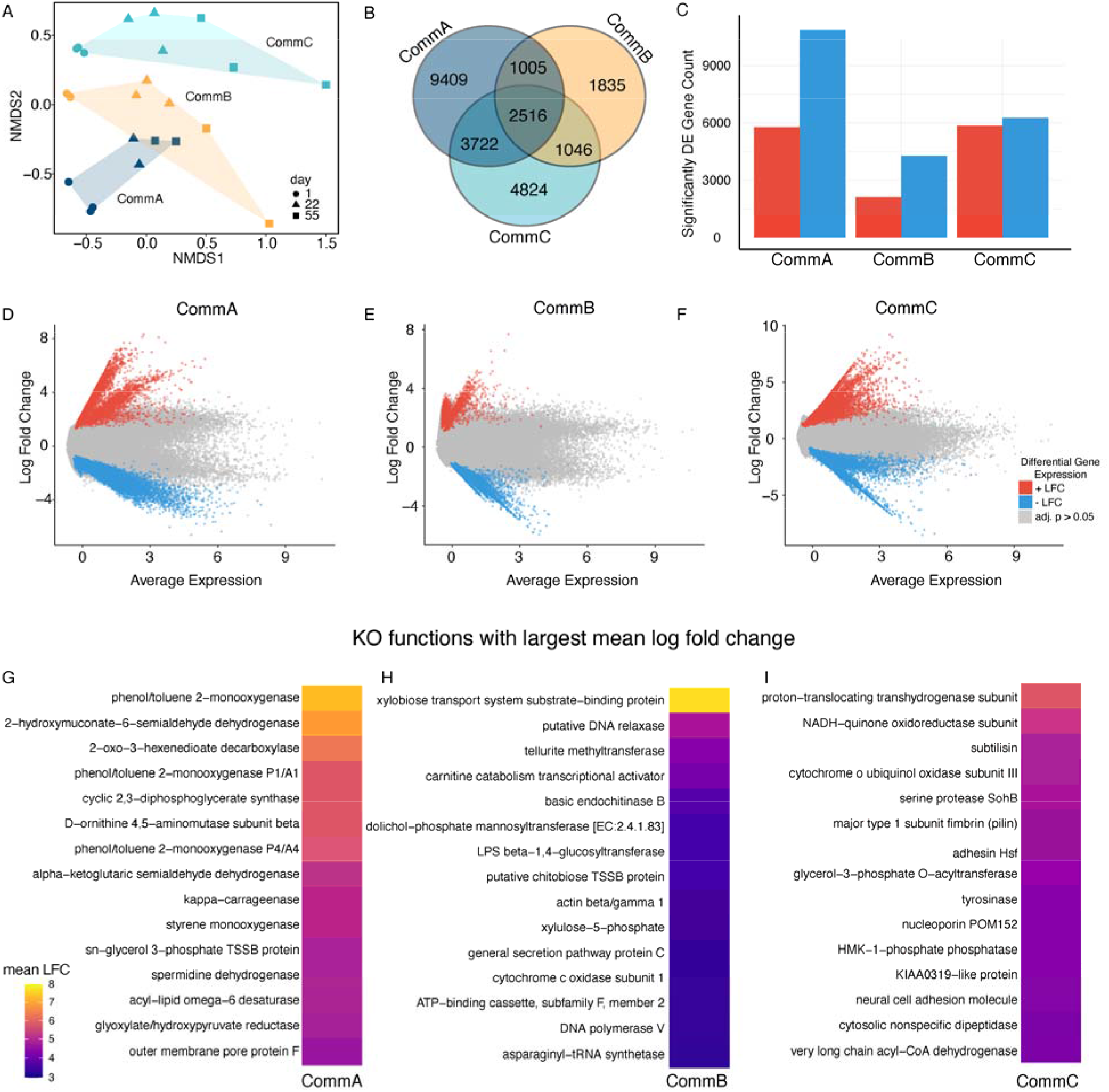
Treatment communities had distinct patterns of gene expression. (**A**) Transcript-based NMDS calculated using Bray-Curtis dissimilarities for all samples within each of the three treatments at days 1, 22, and 55, stress=0.099, k=2 (PERMANOVA, R^2^=0.0.20, F_2,22_=4.46, p = 0.001). (**B**) Venn diagram showing the shared and unique differentially abundant transcripts among the three bacterial communities. (**C**) Bar plot displaying the count of differentially expressed (DE) genes (up regulated in red and down-regulated in blue) between the three bacteria community treatments, highlighting significant differences in gene expression between each community based on community contrasts in our differential expression model. (**D, E, F**) The mean abundance (MA) plots depict the log fold change (LFC) versus the predicted average expression of transcripts for each treatment based on our differential expression model, (**D**) CommA, (**E**) CommB, (**F**) and CommC, (red points are positive LFC, blue are negative LFC, gray are adjusted p-value > 0.05). (**G, H, I**) Heat maps illustrating the 20 most abundant KEGG KO functions, based on the differentially expressed genes averaged within each KEGG Orthology (KO) function for each treatment, (**G**) CommA, (**H**) CommB, and (**I**) CommC. The KOs were ranked from high to low mean log fold change.

Each treatment had unique and shared transcripts (Fig. 5B). Comparison of differential expression between treatments using pairwise contrasts showed 5,767 transcripts with positive log fold change for CommA, and 10,885 with negative log fold change for CommA relative to the mean of CommB and CommC. For CommB, 2,124 transcripts had positive log fold change, and 4,278 had negative log fold change. CommC had 5,851 transcripts with positive log fold change, and 6,257 had negative log fold change, relative to the mean of CommA and CommB. To get an agnostic view of functional expression between treatments, we plotted the model estimate of mean average expression and log fold change for each transcript in each treatment contrast, and color coded according to p-value (Fig. 5D, 5E, 5F). We averaged the log fold change for transcripts mapping to the same KEGG function and ranked them from high to low in order to find the most differentially expressed functions in each treatment (Fig. 5G, 5H, 5I).

In CommA, we found high expression of functions related to phenol/toluene 2-monooxygenase, aminomuconate-semi aldehyde, and 2-oxo-3-hexenedioate decarboxylase, related to aromatic compound degradation (Fig. 5G). For CommB, we found high expression of functions for xylobiose transport system substrate-binding protein, basic endochitinase B, and fructose-6- phosphate phosphoketolase (Fig. 5H). We found support for our physiological measures of high chitinase activity (Figs. 1A and 6A), with basic endochitinase B being one of the top 5 highly expressed functions for CommB. For CommC, there was high expression of functions related to proton-translocating transhydrogenase, glycerol-3-phosphate O-acyl transferase, and very long chain acyl-CoA dehydrogenase, related to fatty acid metabolism (Fig. 5I).

**Figure 6.**
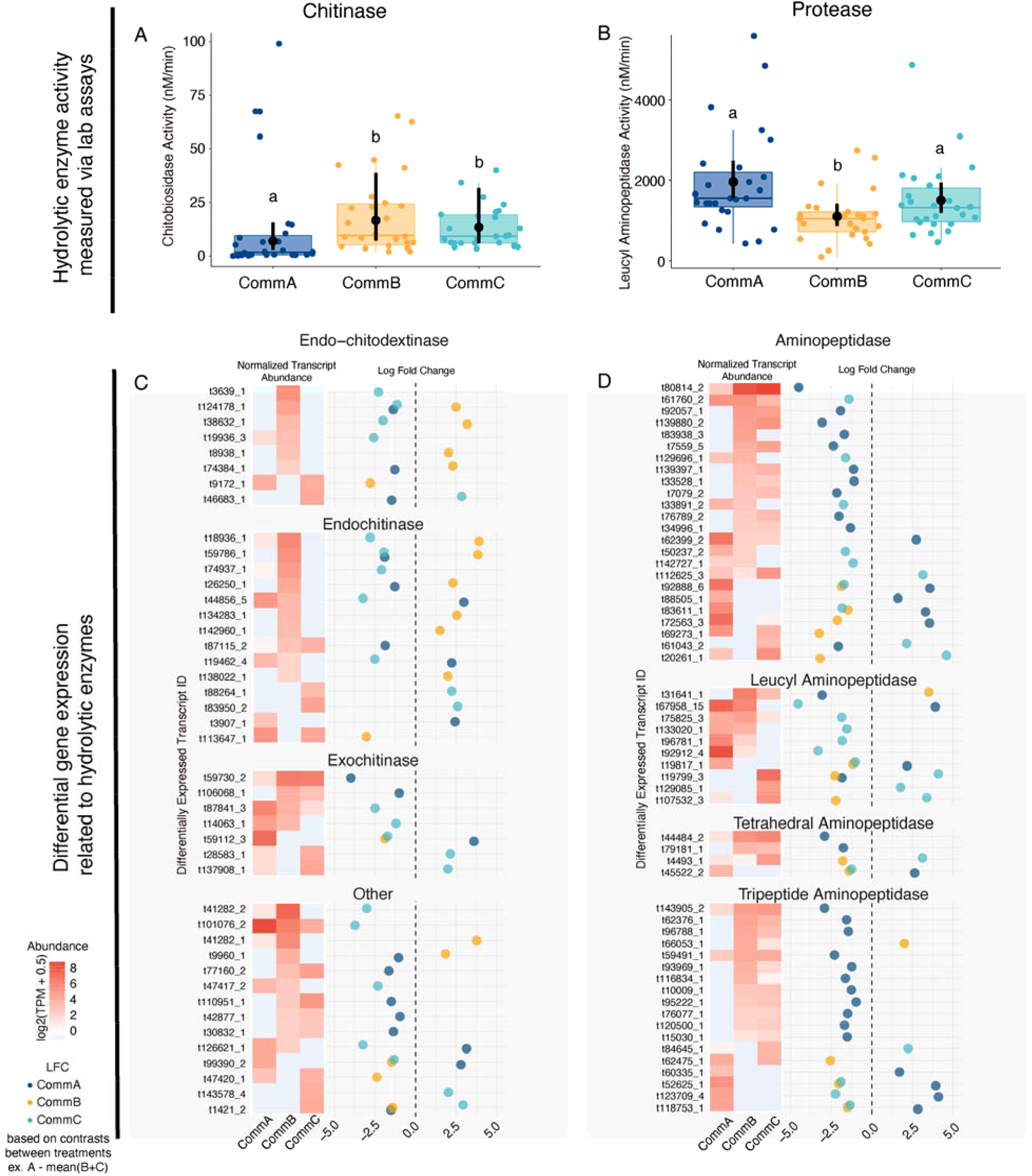
Functional assays of hydrolytic enzyme activity generally matched well with transcript levels of genes related to those functions. (**A, B**) Enzyme activity measured in communities from nine pitcher plants whose samples also underwent downstream metatranscriptomic analysis (within a treatment, three plants x 9 weeks, n=27). Box plots and colored points represent the raw data, with the horizontal bar representing the median values and whiskers showing the 1.5 times the interquartile range. The black points represent the median marginal effects from the model testing the effect of treatment enzyme activity with week as a random intercept, the black vertical bars represent the 95% credibility intervals around each estimate. (**C, D**) Chitinase and protease related transcript comparisons: Within each panel, log fold change (LFC, right) and normalized transcript abundance (TPM, left) of transcripts associated with chitin metabolism KOs (**C**) and protein metabolism KOs (**D**) determined from significantly differential transcript abundances. The log fold change points are colored by treatment and represent the effect size from the differential expression model. The absence of a point for a treatment indicates that the transcript is not differentially expressed for that contrast.

### Linking bacterial physiology with hydrolytic enzyme functional expression

Beyond the general functional differences identified by differential transcription, specific functions related to hydrolysis were examined. We measured chitinase as chitobiosidase and protease as L-amino-peptidase enzyme activities, analyzing only the 9 pitchers which also had metatranscriptomics (3 of 10 plants per treatment). Over 8 weeks, CommA had lower chitinase activity, compared to CommB and CommC (Fig. 6A, Fig. S6). This result was similar to physiological assays in the bacterial communities prior to inoculation (Fig. 1A), but in this smaller subset across time, CommB and CommC were not statistically different. Protease activity in the 9 pitchers was consistent with community assays prior to inoculation, with CommB showing lower protease activity than CommA and C (Fig. 6B, Fig. S6).

We used metatranscriptomic data to explore hydrolysis functions in two ways. First, we investigated the transcript abundances for differentially expressed genes that mapped to chitinase or protease functional KOs. Second, we paired these abundance measures with the log fold change predicted by our differential expression (DE) model. Within this model, contrasts were created to compare the expression levels of each community treatment against the average of the other two (see Fig. 6 legend and methods). This allowed us to identify genes that were differentially expressed in one treatment relative to the others. In Figure 6C, we show all significant DE genes mapping to chitin degradation or transport KOs, including endochitodextinases, endochitinases, exochitinases, and chitin transport systems and putative chitinases. CommB had higher transcript abundances of these chitin related genes, including 12 genes with higher expression in CommB, compared to only six chitin-related genes with higher expression in CommA, and only seven in CommC. Similar patterns emerged with normalized transcript abundance for these same genes (log transformed transcripts per million) (Fig. 6C, left side). In Figure 6D, we show many significant DE genes mapping to protein degradation, including general aminopeptidases, leucyl aminopeptidases, tetrahedral aminopeptidases, and tripeptide aminipeptidases. CommB had lower expression of genes related to protease enzymes, two genes with higher expression in CommB, compared to 12 protease related genes with higher expression in CommA, and seven genes for CommC.

### Linking bacterial physiology with carbohydrate and amino acid metabolic functions

Carbon substrate use profiles were significantly different among all pairwise combinations of the three treatments on both day one and day 55 (PERMANOVA pairwise differences, p < 0.05, Fig. S3, Table S1). Out of the 31 carbon substrates tested, the three communities exhibited differential capacity to use nine (Fig. 7A, Fig. S7). CommB had statistically higher use of N-acetyl glucosamine, beta methyl glucoside, cellobiose, and pyruvic acid methyl ester. CommA showed the highest use of Tween 80, serine, and threonine. Meanwhile, CommC exhibited higher use of galactonic acid gamma lactone and glucosaminic acid.

**Figure 7.**
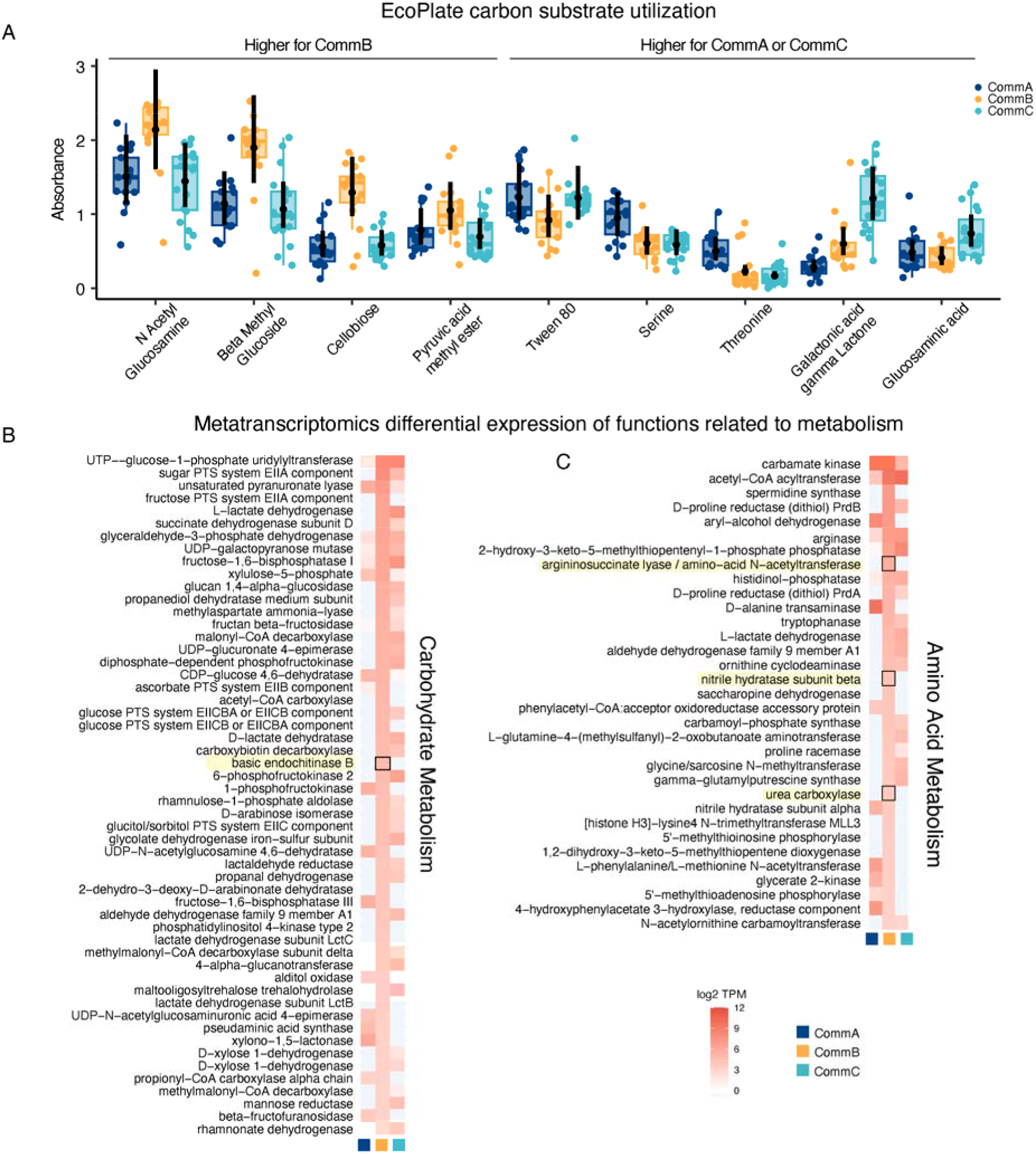
Differences in EcoPlate carbon substrate metabolism among the three treatments. (**A**) A subset of nine of the 31 carbon substrates which showed differences in absorbance among samples of the three treatments. Points are colored by treatment and represent the absorbance for an individual sample for that substrate, day 1 and day 55 samples combined for each substrate. Box plots and colored points represent the raw data, colored whiskers representing data 1.5 times the interquartile range. The black points represent the median marginal effects from the model testing the effect of treatment and substrate on absorbance with week as a random intercept, the black vertical bars represent the 95% credibility intervals around each estimate. (**B, C**) Normalized abundance of KOs associated with (**B**) carbohydrate metabolism and (**C**) amino acid metabolism, using significantly differentially abundant genes summed according to KO function. Heat maps are grouped by treatment and transcripts per million (TPM) are transformed (log2(TPM+0.5)). This subset of KO functions show instances where expression in CommB was significantly higher (by at least 4 log2(TPM+0.5)) than either CommA or CommC. Functions that are particularly relevant for this study are highlighted in yellow.

Relating carbon substrate use from EcoPlate functional assays, we identified significantly differentially abundant transcripts related to carbohydrate and amino acid metabolism (Fig. 7B,C) and aggregated transcript counts by KO function. Because CommB had the largest positive effect on pitcher biomass, we highlighted functions where CommB exhibited markedly higher expression than either CommA or C by at least log_2_ = 4 transcripts per million (TPM). We identified 54 KOs that were higher in CommB, many of which are related to the metabolism or transport of carbohydrates like fructose, glucose, and chitin. Additionally, we identified 33 KOs that were higher in CommB, including the enzymes urea carboxylase, nitrile hydratase, and argininosuccinate lyase. These are related to N cycling, contributing to the breakdown and recycling of nitrogen-containing compounds (29, 30).

### Connecting community functions to metagenome-assembled genomes

In addition to the 16S amplicon sequencing to identify taxa present, we conducted shotgun metagenomic sequencing of 18 samples (3 plants per treatment at day 1 and day 55). The resulting data were used to quantify taxonomy and build high-quality metagenome-assembled genomes (MAGs), which were then connected to the metatranscriptomics data, focusing on hydrolytic enzyme functions. Taxonomic identifications of MAGs were similar to the 16S amplicon results, with some differences likely due to different databases queried for the two data types (Fig. S8). We built 36 MAGs, of which 26 were high quality, eight were medium quality, and two were discarded due to redundancy >10% (Table S4). We calculated the relative abundance of each MAG in each treatment community, mapped the functional transcripts to the MAGs, and selected the KOs associated with chitinase or aminopeptidase (proteinase) activity (Fig. 8). In Figure 8, each bubble represents the number of unique MAG contigs that mapped to each KO, identifying 13 chitin metabolism KOs and 20 aminopeptidase-related KOs. MAG_09 (*Caproiciproducens* sp.) had the highest number of functional chitinase-associated KOs, followed by MAG_41 (*Aquitalea* sp.), and MAG_21, MAG_22, and MAG_23 (Order Sphingobacteriales). Protease activity was well-represented across the MAGs, with the highest abundance of aminopeptidase-related KOs associated with MAG_24 (*Phenylobacterium* sp.), followed by MAG_29 (*Shinella sumatrensis*), MAG_33 (*Comamonas acidovorans*), and MAG_30 (*Sphingomonas* sp.).

**Figure 8.**
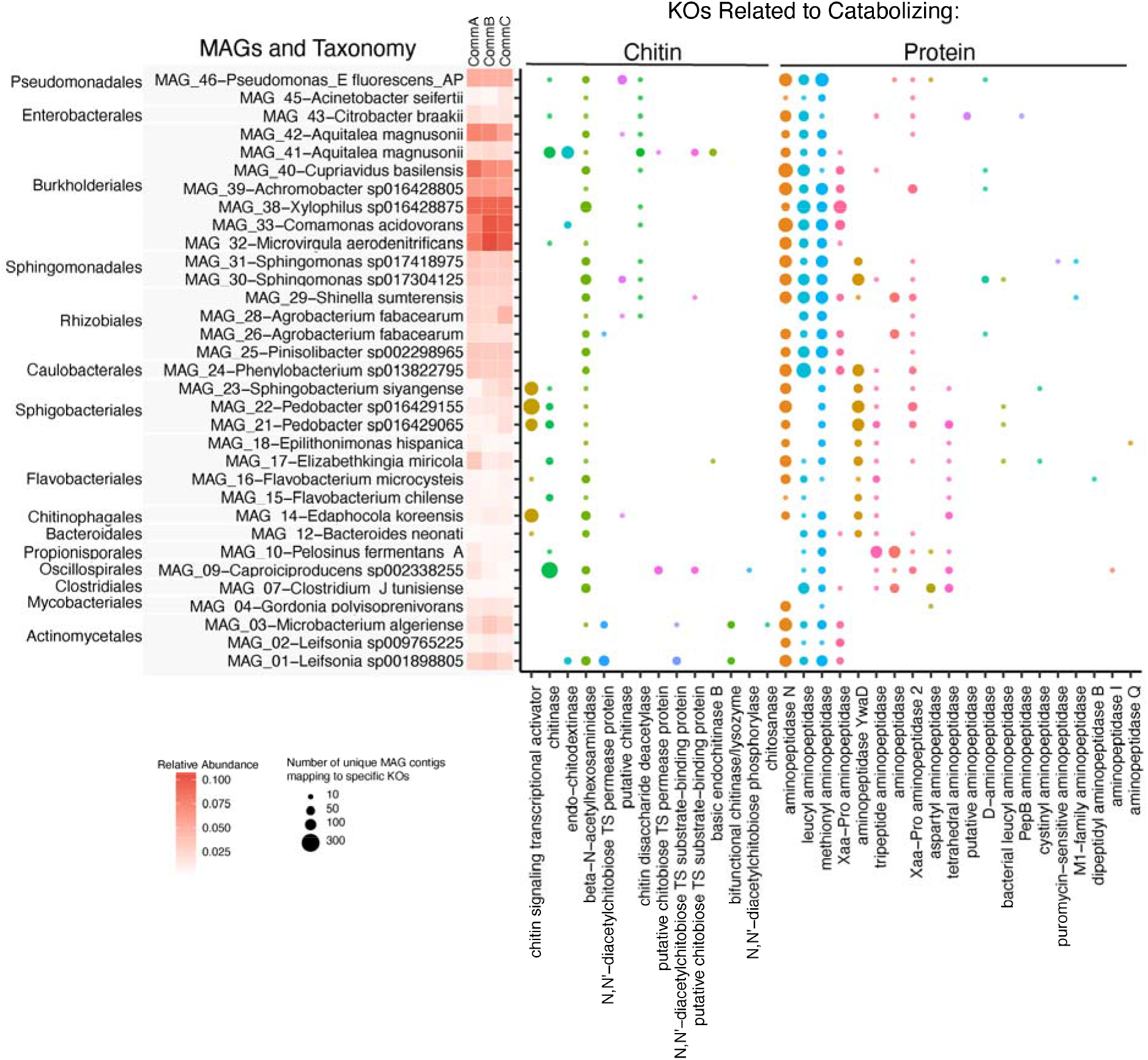
Metagenome assembled genomes (MAGs) enriched in functions related to chitin and protein metabolism. The size of the bubble represents the number of unique MAG contigs mapped to functional transcripts (RNA sequences from the same samples) grouped by KEGG orthology (KO, x-axis) and filtered to include only those KOs associated with targeted hydrolytic enzyme functions (chitin and aminopeptidase metabolizing KOs). The y-axis is ranked by phylogenetic relatedness and shows the MAG identifier along with the associated taxonomy and a heatmap of the relative abundance of each MAG in the three community treatments.

## Discussion

### Bacterial community functioning influences plant traits

Plants and microbes have been associated for approximately 450 million years (31). Over this vast period, many microbes have evolved beneficial interactions with their hosts; however, our knowledge of the mechanisms behind mutualistic effects is still limited. In this study, bacterial community function significantly influenced host growth and nutrient acquisition in the pitchers of carnivorous pitcher plants, almost doubling pitcher biomass, and we identified potential functional drivers of these differences. These results addressed our initial question of whether functionally-different bacterial communities have varying effects on host plant traits, where we found positive effects of CommB on pitcher biomass, length, and carbon content. Functional differences among the bacterial community treatments, such as chitinase activity, were correlated with pitcher growth and plant nutrient acquisition. Many studies provide evidence for how the metabolites produced by microbes incite benefits to their host plants, for example, plant growth promoting hormones (17, 32, 33). However, this is the first example to our knowledge, connecting collective microbial community function to host health using metabarcoding, metagenomics, and metatranscriptomics.

This research leveraged a unique model system, with microbes hosted in aquatic microecosystems inside pitcher-shaped leaves, in contrast to classic non-aquatic phyllosphere systems . One might expect that the aquatic pools would have a lower impact on host health than more intimate associations such as root and leaf associated microbial communities.

However, the positive effect of bacterial communities on leaf biomass *in planta* in just over eight weeks demonstrates a rapid response to their microbial communities for these slow-growing, long-lived plants. In this experiment, a single pitcher was responsible for nitrogen acquisition for the whole plant, so the other pitchers in the experimental plants were filled with sterile water so could not access N. It is likely that nitrogen was translocated out of the target pitcher to support other pitchers (8, 23), reducing the sensitivity or size of the effect on just the measured target pitchers. These plants generally have low nitrogen content compared to other plants (34), suggesting that even small increases in nitrogen could significantly improve their growth and fitness (34). Additionally, several plant traits were examined, and we found that the percent of carbon and nitrogen in the target pitcher was highly correlated with the second- newest pitcher, which was not inoculated, indicating that nutrient translocation was likely occurring. Our findings demonstrate that functionally different microbial communities can influence host plant health, most likely via the release and supply of nutrients that enhance plant growth and fitness.

### Relationship between bacterial composition and function

Recent studies have linked microbially mediated plant benefits to specific taxa. For example, in an experiment with synthetic microbial communities, pathogen suppression in *Arabidopsis* was driven by three specific bacterial taxa (37). In contrast, other studies have observed a community-level effect, such as the positive influence of microbial communities, but not individual strains, on duckweed growth (6, 9, 35). Our study supports the latter situation, as we did not see the presence or absence of specific taxa driving main effects. One possible explanation could be the importance of microbe-microbe interactions, metabolic collaboration in microbial communities is common, making the effects of single taxa less likely (6, 11, 38).

Addressing our second question, although we found clear taxonomic and functional differences in our bacterial communities, most ASVs and MAGs were shared across all three communities and the differences were in rare taxa, without specific taxa appearing to drive primary functional differences. In our study, observed differences in hydrolytic enzyme activity and substrate use were likely affected by small differences in relative abundances of functionally important taxa across our communities. Our results indicate that bacterial community functions are not solely determined by composition but, instead, specific combinations of taxa and differential gene expression lead to distinct functional profiles. Corroborating our findings, a 2023 study with wild pitcher plants found significant functional differences in the microbial community with modest microbial compositional changes following an experimental manipulation, suggesting that differential gene expression, not composition, drove functional change (38). Similarly, in lake microbial communities, functional differences were only weakly related to taxonomic composition (39). Plants often host high bacterial biodiversity, providing complex habitats and resources that create bacterial niches (40). In pitcher plants, the colonization of pitchers by microbial communities are influenced by factors like season, host characteristics, and nutrient inputs, and these differences in microbial community composition likely have important implications of community function (26, 41, 42). In this study, we demonstrated that relatively small differences in bacterial composition and abundance within pitchers can have significant consequences for nutrient cycling from captured prey supplied to the host plant.

### Coupling measured bacterial functions with differential gene expression

Our agnostic approach, to examine total gene expression, allowed us to first examine general functional differences across the three bacterial communities. While substantial differences in hydrolytic enzyme activity were observed between bacterial communities after some time *in planta*, these did not fully explain the observed benefits to plant biomass and pitcher nutrient acquisition, suggesting a more synergistic, collective community effect rather than the additive effect of transcription of selected genes. Each community exhibited differential transcription of a large number of unique genes, many of which are involved in producing enzymes involved in breaking down complex substrates. Interestingly, despite this untargeted approach, CommB had two chitinase-related KOs among its highest transcribed functions, further supporting this as an important function for liberating nutrients including nitrogen from chitin (43).

Additionally, many of the most highly transcribed genes found in CommB mapped to KOs relating to breaking down xylobiose, carnitine, and chitin, indicating this community is efficient at metabolizing complex organic nutrients with probable benefits to the host plant and other members of the microbial community. The catabolism products of these substrates have been found to trigger plant responses, including defense and growth (44, 45). Lastly, while not in the top 20 KO functions, the ACC deaminase gene was significantly higher expressed in CommB. This enzyme plays an influential role in reducing ethylene mediated stress, which can have cascading benefits on growth and development (32). Together, these findings suggest that differential gene expression between microbial communities, due to subtle differences in composition, can contribute to measurable differences in host plant growth.

### Targeted bacterial functions influence plant traits

Few studies have successfully linked direct enzyme activity with transcriptomics for whole microbial communities. Bacterial communities play pivotal roles in nutrient transformations, so in order to better understand community functional dynamics, we linked bacterial physiology with hydrolytic enzyme expression to investigate how chitinase and protease activities *in planta* related to differential gene expression for these functions. CommB and C exhibited higher chitinase activity in physiological assays, and CommB also showed significantly higher expression of genes related to chitin degradation in our metatranscriptomic analyses, connecting active enzyme activity to gene expression. In contrast, CommB had lower leucyl aminopeptidase activity measured via a laboratory assay, while differential gene expression identified many genes mapping to various aminopeptidase enzymes expressed in CommB. These results provide evidence that while culture-based physiological assays are valuable, they fail to completely characterize total nutrient cycling activities present in a community. Pairing physiological assays with metatranscriptomics built a clearer picture of the functional activities present in our communities.

The analysis of carbohydrate and amino acid metabolism across the three bacterial communities supports our findings on hydrolytic enzyme activities, revealing distinct organic substrate preferences among the communities. CommB exhibited higher utilization for substrates like N-acetyl glucosamine (GlcNAc) and cellobiose. GlcNAc is a monomer of chitin, chitobiose, or chitotriose and is used for many critical metabolic functions within bacterial cells (46–48), while cellobiose is a metabolic precursor to glucose and interacts with ?-glucosidase to yield two glucose molecules used in glycolysis (49). Increased transcription of genes related to metabolizing these substrates provides further evidence that CommB is efficiently cycling diverse nutrients in this community, products which are likely important for not only community cross feeding but for nutrient supply to the host plant. Although it has long been assumed that bacterial community enzyme activity is a key mechanism in pitcher plant nutrient acquisition(23, 50), our results are the first to directly measure and provide evidence for this mechanism.

## Conclusion

In this controlled study, plant growth was enhanced by the collective function of a specific bacterial community. The most growth was observed in pitchers hosting a bacterial community with high chitinase activity and high differential expression of genes related to nutrient cycling, supporting the release of nutrients for uptake by the host plant. We characterized collective community composition and function using a suite of complementary approaches that highlighted the complexity and dynamics of plant-microbe interactions and the role microbial communities can play in host health and growth. The benefits of plant-microbe interactions to host productivity are likely often not driven by a single functional parameter, nor a single taxon expressing a gene encoding a specifically useful function. This bacterial community analysis suggests that measurable functional traits can identify relevant bacterial effects in phyllosphere communities. Identification of these relationships contribute to increased knowledge of the effects of bacterial communities on plant traits and can inform other systems with applications in agriculture, conservation, and restoration.

## Methods Experimental design

Mature *Sarracenia purpurea* greenhouse plants (donated by T. Miller, FSU) were surface sterilized and repotted into autoclaved media (7 parts milled sphagnum peat: 3 parts sand) and bleach-sterilized pots. Plant surface sterilization consisted of washing the leaves and roots in distilled water prior to soaking in 1% hydrogen peroxide for 15 minutes (repeated 3X). Plants were transferred into clean plant growth chambers (Hoffman Mfg. Inc. Corvallis, OR USA, SG2- 22) and allowed to acclimatize for 12 weeks prior to the start of the experiment. Care was taken to keep the plants, pots, and all contact with the growth chambers as sterile as possible, however chambers were supplied with f ambient unfiltered air through their vents and when the chamber doors were open. The chambers were configured with a diel light and temperature cycle, 14-hour photoperiod (Photosynthetic photon flux density (PPFD)= 62 μmol m^-2^ s^-1^) at 31 °C and 10 hours dark at 17 °C, with relative humidity at 70%. Additional temperature probes were added to each chamber to monitor chamber conditions.

The three distinct bacterial communities (CommA, CommB, and CommC) used for inoculation were previously collected from wild *S. purpurea* pitcher plants (51). These communities were selected from the ten microcosm communities used by Bittleston et al. 2018 (CommA=M01, CommB=M06, CommC=M09). All ten communities from that study were reconstituted from cryogenic culture and grown in the lab using acidified cricket media (ACM) with 1:1 serial transfers into fresh media every 72 hours for several months. Cricket media was used to establish the bacterial communities and was prepared by acidifying (pH=5.6, 1.0M HCl) ground food-grade house cricket media (3g/L H_2_O Thailand Unique Inc.), then autoclaving. The three communities (CommA, CommB, CommC) were selected based on their differences in chitinase and protease activity (Fig. 2). In addition to three experimental bacterial communities, two experimental controls were ACM only and sterile deionized water only. Both controls contained no bacterial cultures, the ACM control was used to test effects of nutrient media without any bacteria on plants. The water control examined the effect of sterile ACM media with no bacteria.

Ten individual pitcher plants were randomly assigned to each bacterial treatment group and six to each control group, for a total of 42 individual plants. On day 0, on each plant, one new, unopened pitcher was manually opened using sterile gloved hands and carefully inoculated with a bacterial community culture or control treatments until each pitcher was 3/4ths full. For example, if a pitcher held 8 mL when 3/4ths full, it received 8 mL sterile ACM and 0.8 mL bacterial community culture. Due to varying pitcher sizes, some smaller pitchers received lower volumes in the same media:culture ratio. The single pitcher on each plant that received one of the five treatments was the target pitcher (pitcher 1). All other pitchers on each plant received sterile water for the duration of the experiment. For some plant trait measurements, data were also collected on the second-newest pitcher (pitcher 2). We found no statistical differences in the optical density or the bacterial cell concentration (from CFU counts) between our three cultures prior to inoculation.

Because plants did not all have new, unopened pitchers on the same week, new plants were added to the experiment in stages, once a week for 5 weeks until all plants had been inoculated. To ensure there was no effect of this staggered addition of plants, plants were assigned to treatment groups randomly and sample analysis was based on time after inoculation, not date. The cultures for the three bacterial communities were maintained (as noted above) in the lab and added each week to the appropriate plants. There was no change in bacterial community composition in these cultures, and we found that community composition of samples within the same treatment, but added on different weeks, had very tight clustering 7 days after inoculation, regardless of date added. Every 2 days, distilled water was added to the trays under the pots, to prevent splashing and contamination into pitchers. Each week, the plants were rearranged within the chamber and rotated according to a randomization map created using a random number generator.

### Sample collection

Samples were collected weekly, starting the day after inoculation (day 1) 4.0 mL was collected from each leaf using sterile technique. This sample was then aliquoted, 1.5 mL was frozen in sterile screw cap tubes for amplicon sequencing, 750 μL was frozen in cryotubes in liquid nitrogen for metagenomic/metatranscriptomic analysis, 750 μL was mixed with 750 μL 80% sterile glycerol and frozen in screw cap tubes for culture preservation. All preserved samples were stored at -80 °C until nucleic acid extraction. The remaining 1.0 mL was used in the microbial physiological assays described below. The full 4.0 mL was not removed from very small pitchers on day 7 due to limited volumes, but in all cases pitcher size increased very quickly in the first week and full samples were collected at future samplings. In all plants, sample volume (4.0 mL) was replaced with sterile cricket media as a weekly food source, and an additional 2 mL sterile water added to pitcher 1 to replace evaporated fluid (total added liquid each week to all plants = 6.0 mL). There was some damage that occurred to some pitchers during the course of the experiment, plants 25 (CommB), 44 (CommB), 51 (CommC), 9 (CommC), and 29 (CommA) were removed from all analyses.

### Bacterial Physiological Assays

Chitinase and protease activities were quantified weekly on a sample from each treatment pitcher using fluorometric assays in black 96-well microplates (24, 52). Fluorescence emission was measured every five minutes for one hour using a microplate reader (Biotek Synergy Mx Multi-Mode Microplate Reader). Of three types of chitinases found in the *S. purpurea* pitchers communities, chitobiosidase activity was selected for quantification because preliminary data showed higher activity potential than β-N-acetylglucosaminidase and triacetyl chitotriosidase. Chitobiosidase (hereafter chitinase) activity was measured using 200 µL of sample (culture or pitcher fluid) and 50 µL of substrate (0.86 mM 4-Methylumbelliferyl N, N?-diacetyl-β-D-chitobioside in 50 mM Tris-HCl pH 8.0 with 0.1% bovine serum albumin) as described in (24).

Standards of 4-methylumbelliferone were made from 0 to 1 µM concentrations and read at 360 nm excitation and 420 nm emission. L-amino peptidase (protease) activity was measured using 50 µL of sample added to 25 µL of 50 mM Tris-HCl pH 7.5 and 75 µL of substrate (2 mM L-leucine-7-amido-4-methylcoumarin hydrochloride in nanopure water). Standards of 7-amino-4- methylcoumarin were made from 0 to 1000 nM concentrations and read at 380 nm excitation and 460 nm emission. Using the standard curves for each assay, fluorometric readings were converted to product concentrations and plotted through time to get the linear enzymatic rate for each sample (24).

The “functional fingerprint” for each bacterial community was measured by examining the community’s capacity to use 31 carbon substrates. These measurements were conducted on the first and last day of the experiment (day 1 and 55) using Biolog EcoPlates™. EcoPlates were inoculated with 100 µL of diluted sample (1:40 in sterile DI water) and allowed to incubate in the dark at room temperature, absorbance readings were collected every 24 hours until a maximum color change had been observed (72 hours). The changes in color were quantified by reading the absorbance using a plate reader (above) at a wavelength of 590 nm. Water was used as a control blank and subtracted from each absorbance reading prior to analysis.

Experimental controls were removed from the EcoPlate analysis. After an initial analysis, extreme outliers at day 55 were removed the EcoPlate analysis, including plants 34 and 15 from CommA, 27 from CommB, and 28, 21, and 41 from CommC.

Bacterial community growth metrics (maximum growth rate, doubling time, carrying capacity) were measured using optical density at 590 nm for bacterial communities prior to inoculation (Day 0) and from pitcher fluid samples at the end of the experiment (day 55). Bacterial cultures and pitcher fluid samples (2 µL) were added to 198 μL R2A media in a clear flat bottom 96 well plate (53). Plates were incubated in a plate reader (Tecan Spark) at 18 °C and absorbance was measured every 30 minutes for 48 hours. Bacterial growth metrics were analyzed using the R growthcurver package (54). The microbial community respiration was measured (as CO_2_ produced) for both the bacterial cultures prior to the experiment (Day 0) and for pitcher fluid samples at the end of the experiment (day 55), using the MicroResp system (55). Samples were added to a sterile deep well plate (250 µL) along with 250 µL acidified cricket media. The plate was fitted with a silicon mat with small holes and an indicator plate, both plates were clamped into the MicroResp apparatus and incubated in the dark at room temperature. The indicator plate contains a pH sensitive dye (cresol red 12.5 ppm, wt/wt) that shifts from magenta to yellow in the presence of carbon dioxide. The absorbance at 570 nm was measured prior to incubation, and consecutively after 6, 24, and 48 hours. Carbon dioxide concentrations were calculated using a standard curve created by incubating an indicator medium with a series of known carbon dioxide concentrations. Respiration rates were calculated for the first 24 hours of respiration as a change in the CO_2_ produced over the first 24 hours.

### Plant functional traits

Pitcher morphological traits (56), including length, width, and opening aperture (56) were measured weekly for eight weeks (day 0, 7, 14, 21, 28, 35, 42, 49, 55). For each plant, the number of mature leaves was recorded at the beginning and end of the experiment. The length of each pitcher (rhizome to tip) was measured with a flexible tape (cm) and the pitcher diameter (at widest point), and aperture of the pitcher, will be measured with digital calipers (mm). Tools were carefully sanitized with 70% ethanol between each measurement. At the end of eight weeks, two pitchers were cut at the base for biomass measurements, the target pitcher (pitcher one) and a secondary mature pitcher (second-oldest pitcher, pitcher two). Biomass was measured by first taking a wet mass, drying the plant tissue at 65 °C for 72 hours, then recording the dry mass. Plant tissue was homogenized in a spice grinder, for a subset of plants (n= 15), we also harvested the remaining pitchers, roots, and rhizomes to investigate plant level differences in biomass and C: N for plants in different treatment groups. Whole plant (pitcher, root, rhizome) samples were collected for nutrient content analysis at the end of the experiment (three whole plants from each treatment (n=15) and the two main pitchers from the remaining plants). Plant tissue was dried at 65 °C for 72 hours and stored in the dark at room temperature until samples were homogenized. Powdered samples were analyzed using a CN elemental analysis (Flash EA1112), the proportion of carbon and nitrogen for each tissue sample was calculated using the mass of the nutrient divided by the mass of the ground sample.

Photosynthetic rates (µmol CO_2_ m^-2^ s^-1^) of the target pitcher (pitcher one) and secondary mature pitcher (pitcher two) were measured separately at the end of the experiment (day 55) using a LICOR-6400 photosynthesis system (LICOR Inc., Lincoln, NE) and leaf chamber fluorometer (LCF 6400-40). Measurements were taken between 1000 and 1300 h on plants removed from the growth chambers, pitchers were filled with distilled water.

Photosynthetically active radiation (PAR) was set to 1200 µmol photons m^-2^ s^-1^ for 5 min. A subset of plants (three plants from each of the five treatments, n=15) had photosynthetic rates measured on the youngest mature pitcher on each plant, prior to inoculation and addition to the experiment.

The maximum quantum efficiency of photosystem II photochemistry (F_v_/F_m_)(57) was measured once daily for one week, then weekly for 8 weeks. A portable multimode chlorophyll fluorometer (Opti-Sciences) was used to measure the maximum quantum yield (F_v_/F_m_) of chlorophyll fluorescence after a 25-minute period of dark acclimation using plastic leaf clips(34). The fiber optic cable was wiped with 70% ethanol between each plant and the plastic leaf clips were sanitized with 10% bleach, rinsed with DI water, and allowed to air dry after each measurement.

Many plant traits were measured (leaf length, width, aperture, biomass, photosynthetic efficiency, photosynthetic rate, pitcher carbon, pitcher nitrogen, total number of leaves, new leaves). The collinearity in the response variables was assessed using a principal components analysis (PCA) on scaled data, dimension 1 explaining 77.1% of the variance and dimension 2 explaining 7.9% (Fig. S9). Along dimension 1, pitcher morphological measures (length, width, aperture, biomass) were tightly correlated with each other. Pitcher dry biomass was selected as the representative measure of plant size. Pitcher nitrogen and carbon content were selected as measures of nutrient assimilation (Fig. S9).

### 16S rRNA sequencing and analysis

To characterize bacterial community composition, DNA was extracted from all samples (CommA, CommB, CommC, ACM & water controls, on day 8, 15, 22, 29, 36, 43, 50, 55, n=310) using the DNAdvance Genomic DNA Isolation Kit (Beckman Coulter A48705). Samples (750 µL) were bead beaten in lysis buffer at 2400 RPM for 10 min, and then incubated at 55 °C shaking (150 RPM) overnight before proceeding with the extraction (halving all the reagents but following the protocol per the manufacturer’s directions), each 96-well plate included one negative control. DNA was quantified using AccuClear Ultra HS dsDNA kit (Biotium #31028). 16S rRNA gene amplification and library preparation and sequencing was conducted by the Environmental Sample Preparation and Sequencing Facility at Argonne National Laboratory on a 151 bp x 12 bp x 151 bp Illumina MiSeq run targeting the V4 region of 16S rRNA using the 515F and 806R primers (58).

The 16S rRNA gene sequences were processed in QIIME2 (version 2022.8), demultiplexed using no-golay error correction and quality filtered to remove reads with a mean score less than 20, and trimmed to the sequence length of 150 base pairs. The DADA2 (74) module was used to denoise sequences and generate amplicon sequence variants (ASVs) (59, 60). Taxonomy was assigned using the classify-sklearn method which is a Native Bayes classifier, and a pre-trained classifier made with Silva v. 138 database containing 99% amplicon sequence variants (ASVs) from 515F/806R region (61). The phylogenetic tree was built using multiple-sequence alignment via SEPTT and phylogenetic reconstruction via FastTree, both via a QIIME2 plugin (62, 63). Taxonomy and ASV tables were migrated to R for further analysis and visualization using qiime2R, picante, and phyloseq (64–66).

Across 316 samples (310 pitcher fluid samples and 5 negative controls) *DADA2* generated 2633 ASVs. Contaminant ASVs were identified and discarded using the *decontam* R package (67), the *decontam* method “prevalence” was used which identified ASVs based on their presence and abundance in our negative controls. Data was quality filtered to remove non-prokaryotic ASVs and ASVs classified as mitochondria or chloroplasts, and only include observations with at least 10 sequences and samples with at least 1000 sequences, resulting in 310 pitcher fluid samples (day 8 through day 55) with a cumulative 1036 distinct ASVs, with 25317 mean reads per sample (min reads/sample = 1669; max reads/sample= 57927). Samples were rarified down to 1669 reads, resulting in 310 samples with 1036 ASVs. A negative binomial model was built using the *brms* package to investigate the effect of a treatment group on alpha diversity (68).

For ASV-level differential abundance explained by the treatment groups, an analysis of compositions of microbiomes with bias correction (*ANCOMBC2*) was used, this is a Bayesian statistical approach that accounts for sampling fraction, normalizes the read counts, and controls for false discovery rates (fdr) (28). Quality controlled but unrarefied ASV counts were analyzed using the *ANCOMBC* function, adjusted p-value method was set to fdr, prevalence (prv_cut) was set to 0.3, the level of significance (alpha) was set to 0.05, pairwise directional test (pairwise) was set to TRUE, with the rest of the parameters left as default. Significantly differentially abundant ASVs were identified across the treatment groups and through time.

### Metagenomic and metatranscriptomic sequencing and analysis

The pitcher fluid from three of the 10 pitchers in each treatment group were selected from three time points (3 plants x 3 treatments x 3 timepoints (day 1, 22, 55), n=27) for simultaneous DNA/RNA extractions for metagenomic and metatranscriptomic sequencing using the Qiagen AllPrep Bacteria DNA/RNA/Protein Kit (Cat 47054), according to manufacturer directions.

Briefly, flash frozen samples were thawed and pelleted, pellets were resuspended in lysis solution and bead beat for 10 minutes at 2400 RPM after which 350 µL of lysate was transferred to a spin column were the DNA was collected in the column and the flow through was further processed for RNA extraction. Samples were eluted in nuclease free water and stored at -20°C until quantification and library preparation. Nucleic acids were quantified using a fluorometer (Qubit, Invitrogen). Libraries were prepared for the full 27 RNA samples using NEBNext rRNA Depletion Kit (NEB#E7850) and NEBNext Ultra II Directional RNA library prep kit for Illumina (NEB#E7760) using manufacturer protocols. Metagenomic libraries (DNA) were prepared for a subset of these samples (3 plants x 3 treatments x 2 timepoints (day 1, 55), n=18) using NEBNext Ultra II FS DNA library prep kit for Illumina (NEB#78055). Sequencing was done on a NovaSeq partial lane to a sequencing depth of 55 Gb per library at Novogene Co. (Sacramento, CA USA).

Shotgun metagenomic and metatranscriptomic sequences were processed on BSU’s Borah cluster. Adapters and low quality sequences were trimmed using Trimmomatic (0.39) with the following settings: SLIDINGWINDOW:4:20 MINLEN:25 ILLUMINACLIP:TruSeq3-PE.fa:2:40:15 based on the FastQC (v0.12.1) and viewed with MultiQC (v1.6) quality reports (69–71). Metatranscriptomic sequences were further processed to remove ribosomal RNA using the default settings (-t 4, -l 150, -e rrna) in Ribodetector (72). Two co-assemblies were produced (metagenomic and metatranscriptomic) using MEGAHIT (v1.2.9) (73, 74).

For the metagenomic analysis, the “sketch”, “gather”, and “tax annotate” functions within sourmash (version 4.8.2) (75) using kmer=31 were used to assign and quantify taxonomy using the full GTDB database (R08-RS214 403k) (76). Contigs in the coassembly and paired-end reads were binned using MAXBIN2 (2.2.7), CONCOCT (1.1.0), and Metabat2 (2.12.1), consensus bins were produced using DASTool (1.1.6) (77–80). Completeness and contamination of each metagenome assembled genome (MAG) were calculated using the presence of single-copy marker genes with CHECKM2 v.1.0.11 (81). Comparisons and dereplication of MAGs employed dREP v.2.2.1 (82). Taxonomy was assigned to the quality-controlled MAGs using the full GTDB database within sourmash. Bins were quality ranked according to their completeness and contamination (>90% completion, <5% redundancy), medium quality (>50% completion, <10% redundancy), low quality (<50% completion, <10% redundancy), or discarded (>10% redundancy) (83, 84). The abundance of each MAG in each treatment was calculated by first building an index for each MAG using the ‘index’ function in bwa (85), using bwa and samtools (86) to align the individual forward and reverse sample reads to each MAG. The phylogentic tree for the MAGs was built by first extracting the marker genes from each MAG using the ‘ineage_wf’ and ‘qa’ tools from CHECKM2 (81) and then multiple sequence alignment with MAFFT (62). The phylogenetic tree was built with IQ-Tree (87) and parsed in R. Predicted gene sequences from the metatranscriptomic samples were mapped to a custom database built with the MAGs, and then used blastn (88) to quantify the number of contigs in each MAG that matched with each predicted gene.

For metatranscriptomic analyses, we used SeqKit (89) and VSEARCH (90) to look at the quality of coassembly before and after chimera removal. We removed 23 contigs that surpassed the 50,000 base pair threshold. Open reading frames (ORFs) were predicted using Prodigal (v2.6.3) (91). We used CDHIT to remove redundancy in our predicted proteins by clustering transcript. Transcript abundance quantification was done in Kallisto (v1.9.0) and resulted in transcripts per million (TPM) and counts for each transcript and sample (92). KOFAMSCAN (1.3.0) was used to bin ORFs into KOs using the full KEGG database (93).

Differential transcript abundance was quantified using *Dream* within the *variancePartition* package (94) with treatment set as a fixed effect and plant ID set as a random intercept to account for repeated measures across three time points. The dream model was used to estimate weights using our pre-defined linear mixed model and then fit the model using the Satterthwaite approximation. The coefficient table was filtered down to those z-scores greater than or equal to the critical value of 1.645 resulting in those genes that are differentially expressed across our treatments. Within this model, contrasts were created to compare the expression levels of genes in each community treatment against the average of the other two: CommA vs (CommB + CommC)/2, CommB vs (CommA + CommC)/2, CommC vs (CommA + CommB)/2.

### Statistical analyses

All statistical analyses were conducted in R version 4.2.2 (95). Alpha diversity metrics (Hill Number 1) were calculated using the *vegan* package (96) and additional processing used *tidyverse* and *janitor* (97–99). For ASV-level differential abundance explained by the treatment groups, we used *ANCOMBC* (ancombc2(data = tse, assay_name = "counts", tax_level = NULL, fix_formula = "treatment", rand_formula = "(1 | week)", p_adj_method = "fdr", pseudo_sens = TRUE, prv_cut = 0.3, group="treatment", alpha = 0.05, n_cl = 3, verbose = TRUE, global = TRUE, pairwise = TRUE)) (28). Significantly differentially abundant ASVs were visualized with box plots and a heatmap (100) by transforming the normalized reads: (log2(read count+0.5)).

Community similarity in ASVs and EcoPlate carbon substrate use among samples in each treatment group was determined using the vegan package (96). Compositional differences (beta diversity) and changes across the three bacterial community treatments from day 8 to 55 were assessed using phylogenetic methods, unweighted UniFrac (101). Mantel tests (96) assessed correlations between bacterial composition and hydrolytic enzyme activity or pitcher biomass. To visualize the similarity in EcoPlate carbon substrate use among samples and treatments, a *metaMDS* analysis with Bray-Curtis distance at two dimensions (k=2) was utilized. In both cases, non-metric multidimensional scaling (NMDS, (96)) visualized the dissimilarities calculated between each pitcher fluid sample. A non-parametric multivariate analysis of variance test (*PERMANOVA*, *adonis2* function, by=“margin”) was used to assess any differences between treatments, and *pairwise adonis* was used to assess the effects of treatment and time, while accounting for repeated measures for each pitcher (96). Dispersion, or the homogeneity in dispersion between treatments, was calculated using the *betadisper* function (permutations compared using a post-hoc Tukey test, (96).

Metatranscriptomic statistical analyses began with a matrix of filtered genes (length>500bp), samples were normalized using the *calcNormFactors* function in the *variancePartition* package (1.28.9). Differential transcript abundance was quantified using *Dream* within the *variancePartition* package (94). In order to link specific taxa to differentially expressed functions, significantly differentially abundant contigs were mapped to the MAGs using a custom database built from the MAG contigs and *BLAST* (2.14.0) (88).

To isolate the impact of functionally distinct bacterial communities on plant traits (after 8 weeks *in planta*, Questions 1 and 2), we developed Bayesian generalized linear mixed models (GLMMs) using variables based on the causal assumptions of directed acyclic graphs (DAGs). Directed acyclic graphs help identify which observed and unobserved variables to condition upon to remove non-causal associations between variables and isolate mechanisms of interest, under the assumption that the depicted pathways between variables in a research design are accurate (102, 103) Therefore, GLMMs contained a variable set designed to isolate the effects of treatment and enzymatic activity from the other unspecified and unrecorded impacts of our bacterial treatments (Fig. S1) (68). Pitcher biomass was selected to represent the morphometrics and growth, along with leaf nitrogen content representing leaf nutrient acquisition. To consider non-independence of repeated measures of pitchers over time, models included a random intercept for pitcher identity. Other random effects like inoculation date and initial pitcher volume were not accounted for in the model but controlled for by the randomized design.

Bayesian hierarchical models using a gamma (log-link) GLM were fitted using the *brms* package (68, 104) and visualized using *bayesplot*, *ggeffects*, and *tidybayes* (105–107). Bayesian GLMs are robust to small sample size and outliers, and the gamma distribution best fits the data generating processes (108, 109). The continuous predictor variables were standardized and mean centered using the *standardize* package (108). In all models that contained the treatment group as a predictor, we set ACM as the baseline, to compare the effects to pitchers with food but no inoculated bacteria. Default uninformative priors were used, convergence and mixing of chains and unimodality in posterior predictions were visually assessed, and all R-hat values were close to 1.0 (68). The model fit was evaluated using the posterior predictive check function in *brms* (68).

## Supporting information

Supplemental Results

## Acknowledgements

We would like to acknowledge Stacey Pedraza for help with processing samples, the late Bill Freutel and other members of the Earth, Wind, Fire Lab for use of lab space, Dr. Linda Reynard for FLASH assistance, and Franco Nero and Jacob Heil for help with experiments. We offer our gratitude to Dr. Allison Simler-Williamson and the Ecological Statistics community at BSU for valuable feedback. We would like to thank Dr. Ben Baiser and Dr. Zac Freedman for helpful suggestions and feedback on the final manuscript.

## Data Availability Statement

Scripts and data associated with this manuscript are available at: https://github.com/jessibernardin/microbial-function-plant_trait. The raw reads for 16S rRNA amplicon sequencing, raw reads for whole genome shotgun sequencing, whole genome metatranscriptomic sequencing, and nucleotide sequences of MAGs have been deposited in the National Center for Biotechnology Information (NCBI) under the project number PRJNA1028624.

