## Supplemental Results for "Microbial communities increase host plant leaf growth in a pitcher plant experimental system"

Microbial community function increases leaf growth in a pitcher plant experimental system

This PDF file includes:

Supplementary Results Figures S1 to S9

Supplementary Results Tables S1 to S4

Other Supplementary Materials for this manuscript include the following:

Github: https://github.com/jessibernardin/microbial-function-plant_trait.git

Zenodo: https://doi.org/10.5281/zenodo.10055295

NCBI: https://www.ncbi.nlm.nih.gov/bioproject/PRJNA1028624

**Supplementary Figures**

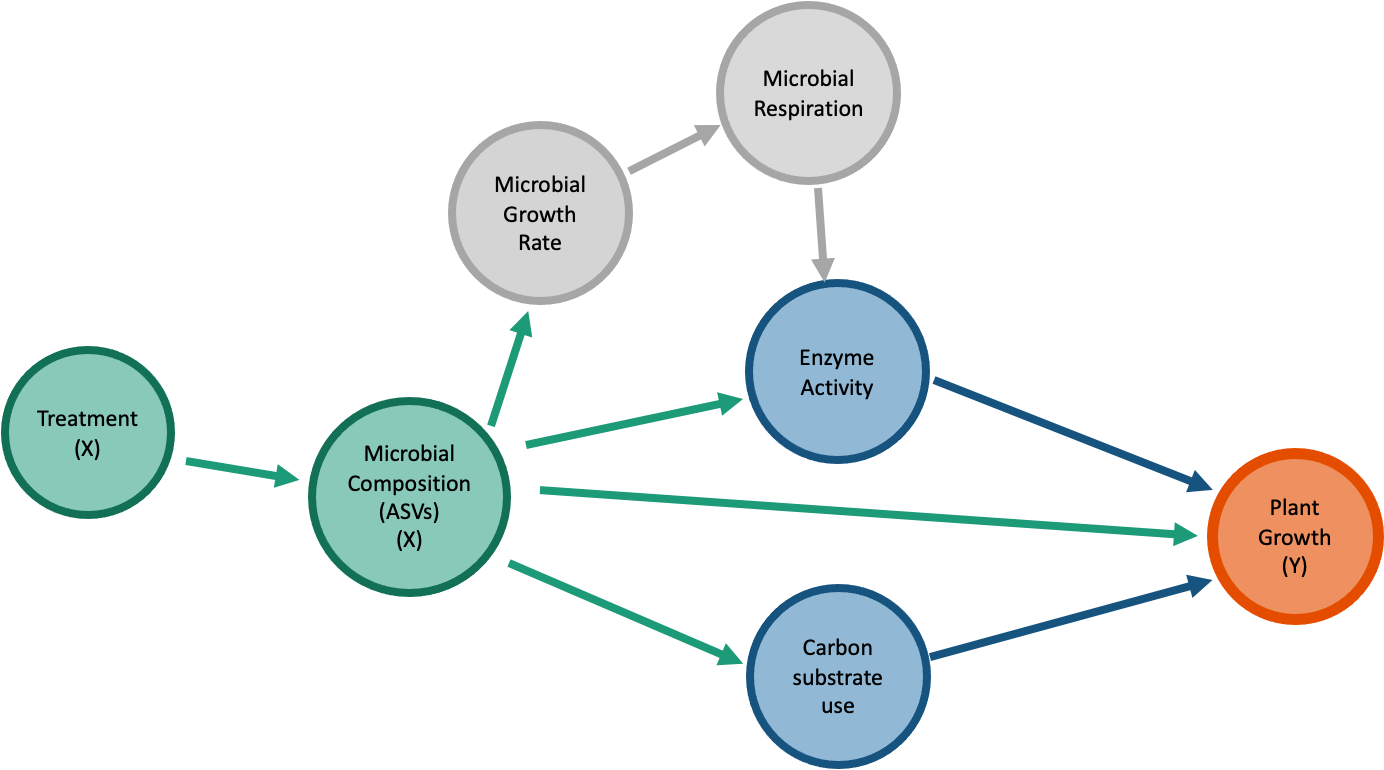

Figure S1. Directed Acyclic Graph (DAG) visualizing the predictor variables (microbial functions and composition) we hypothesize influence plant growth traits (biomass, leaf nitrogen content).

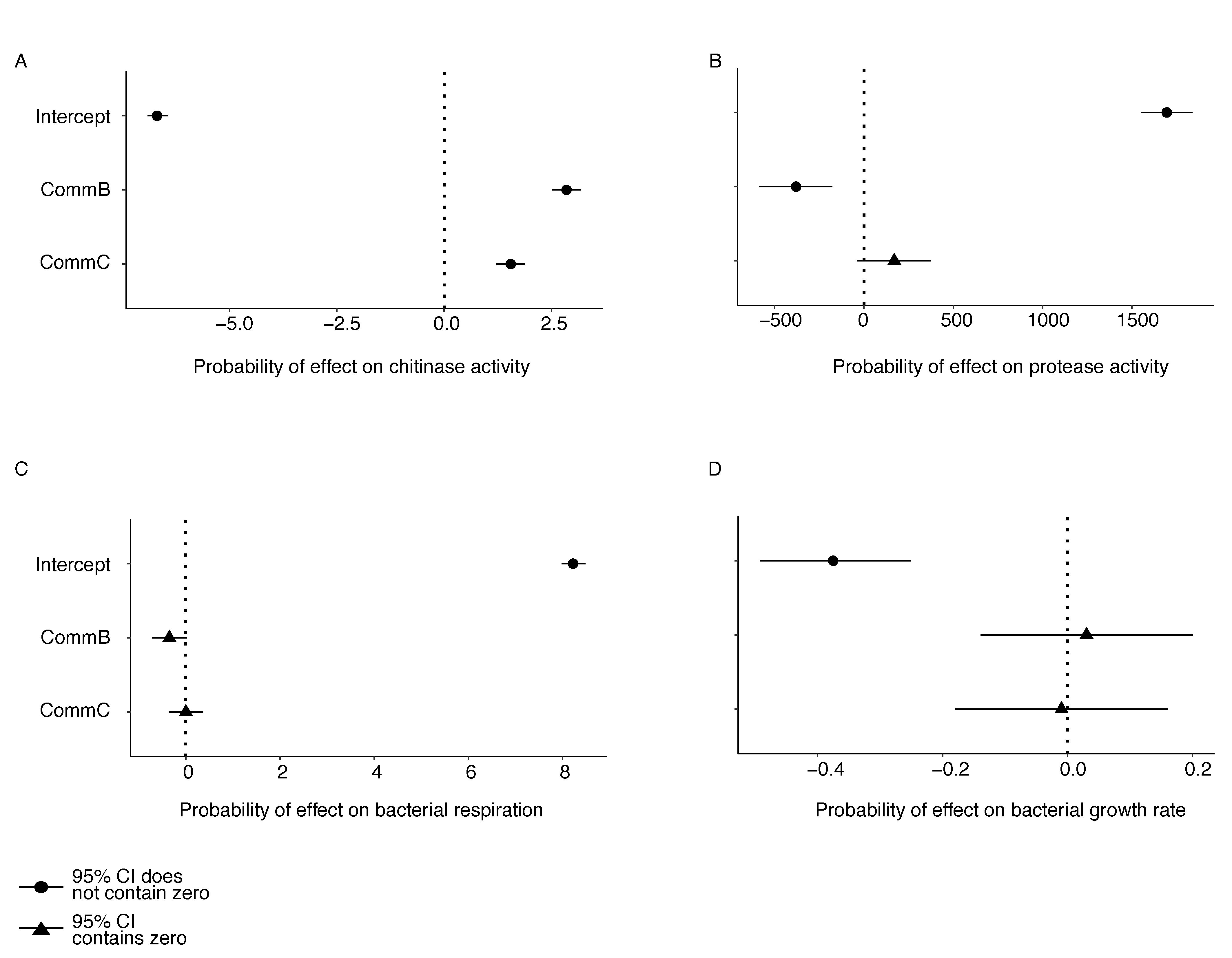

Figure S2. Functional differences in starting bacterial communities (before inoculation). Parameter estimates for the probability of differences between bacterial community treatments on (**A**) chitinase activity, (**B**) protease activity, (**C**) bacterial respiration of bacterial community culture, and (**D**) bacterial growth rate before the start of the experiment. Black points indicate median parameter estimates with associated 95% credibility intervals. Parameters with 95% credibility intervals that did not include zero (dashed vertical line) were considered nonzero effects on the response (circles vs. triangles). CommA is set as the baseline predictor in all models containing treatment.

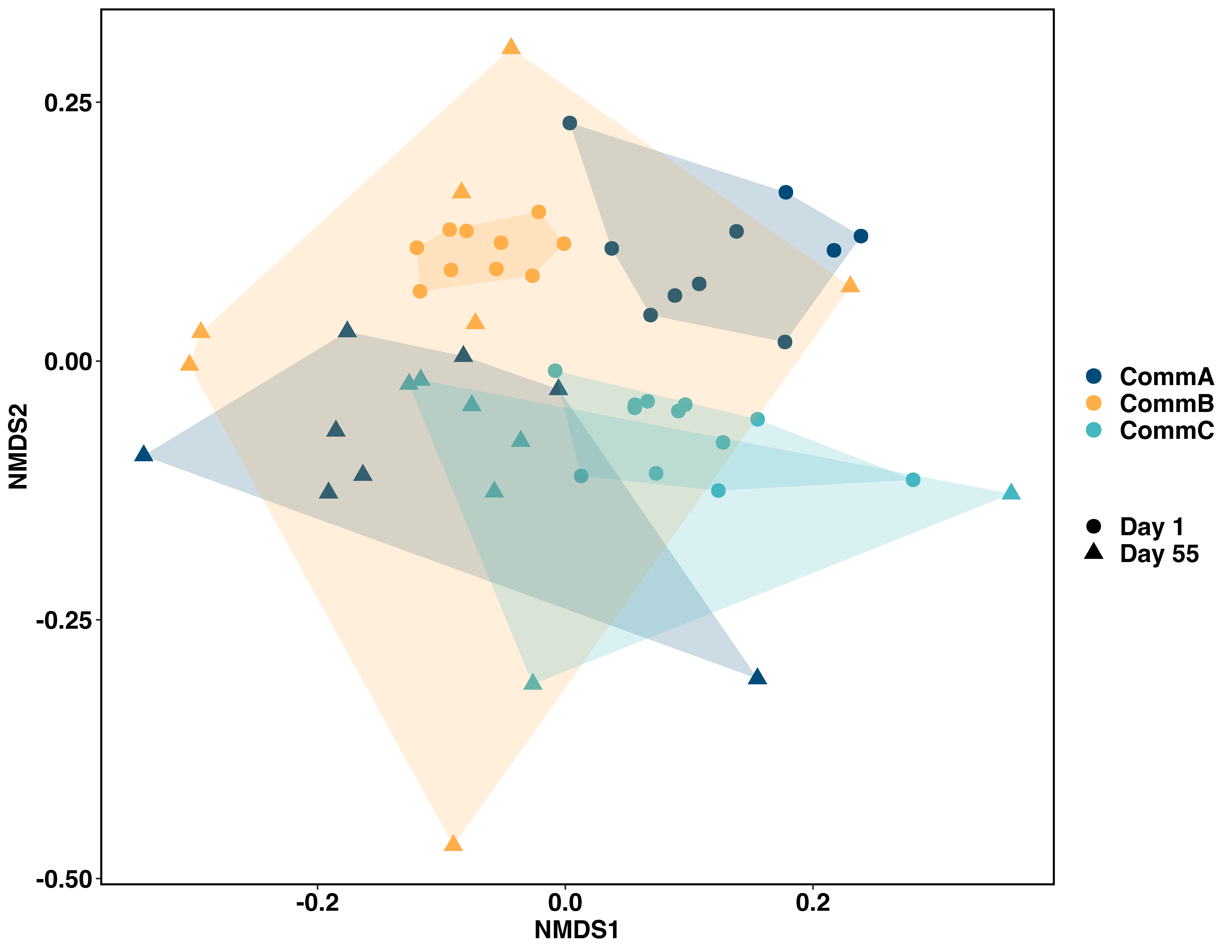

Figure S3. Non-metric multidimensional scaling (NMDS) based on Bray Curtis dissimilarity from Biolog EcoPlate physiological profiles for pitcher bacterial communities at day 1 and day 55. Treatments are coded by color and time by shape. Significant (p<0.05) differences between treatment groups within individual time points and between day 1 and 55 (Table S1).

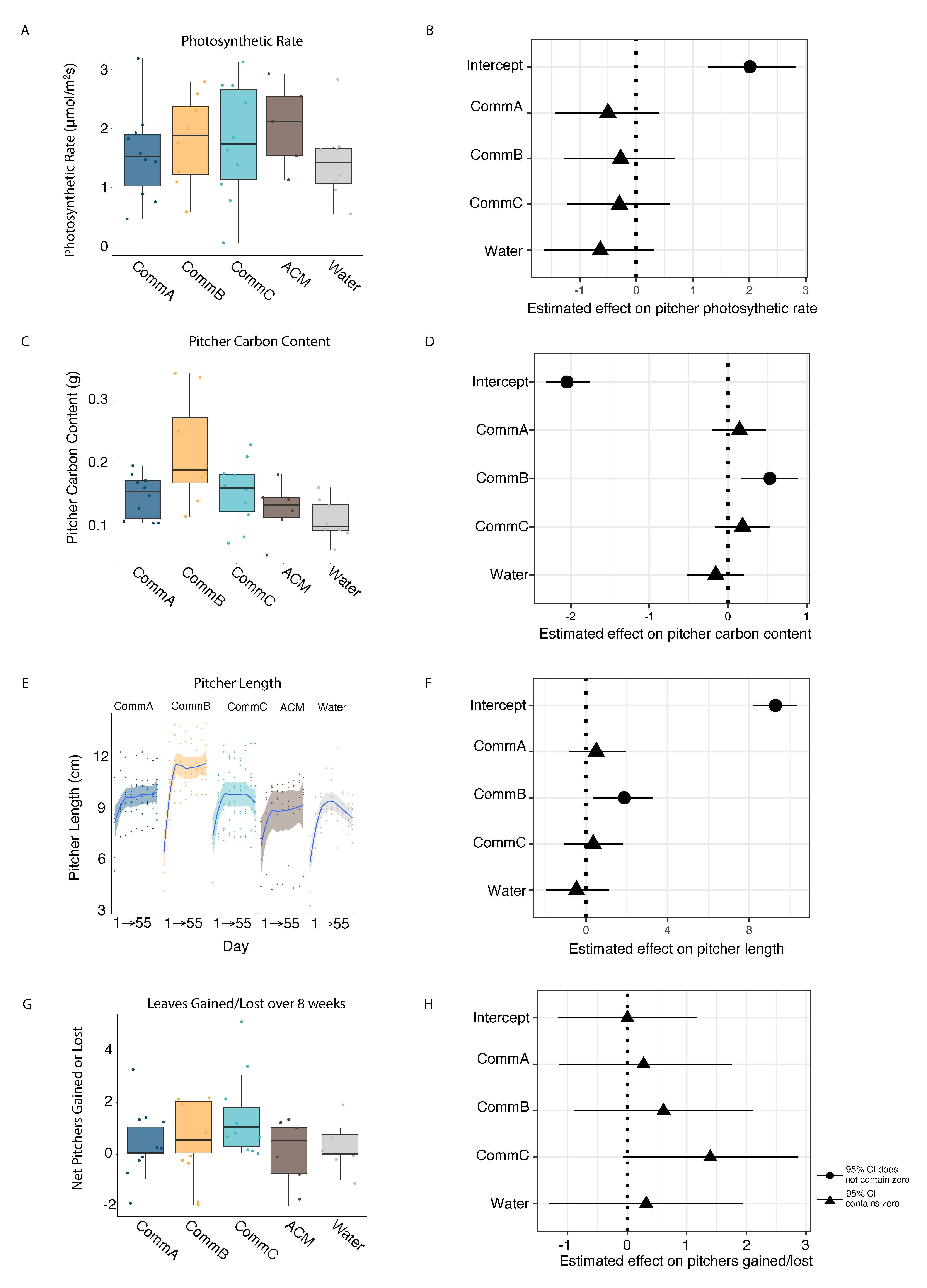

Figure S4. Exploring the effect of treatment on other plant traits. (**A**) The photosynthetic rate (µmol CO_2_ m^-2^ s^-1^) was measured on the treated pitcher at the end of the experiment (day 55). (**B**) Posterior parameter estimates for the effects of treatment on pitcher photosynthetic rate. (**C**) Pitcher carbon content was measured at the end of the experiment; carbon content was higher in CommB compared to the experimental and control treatments. (**D**) Posterior parameter estimates for the effects of treatment on pitcher carbon content. (**E**) Pitcher length was measured weekly for each treated pitcher; pitcher length was higher in CommB compared to the experimental and control treatments and correlated with other plant morphology measures. (**F**) Posterior parameter estimates for the effects of treatment on pitcher length. (**G**) Number of net pitchers gained or lost (starting number of pitchers minus ending number of pitchers) across all treatments. (**H**) Posterior parameter estimates for the effects of treatment on net pitchers gained or lost. **(B, D, F, H)** Symbols represent the median parameter estimates and lines represent the 95% credible intervals for the parameter estimate. Parameters with 95% credibility intervals that did not include zero (dashed vertical line) were considered nonzero effects on the response (circles vs. triangles), ACM is set as the baseline predictor.

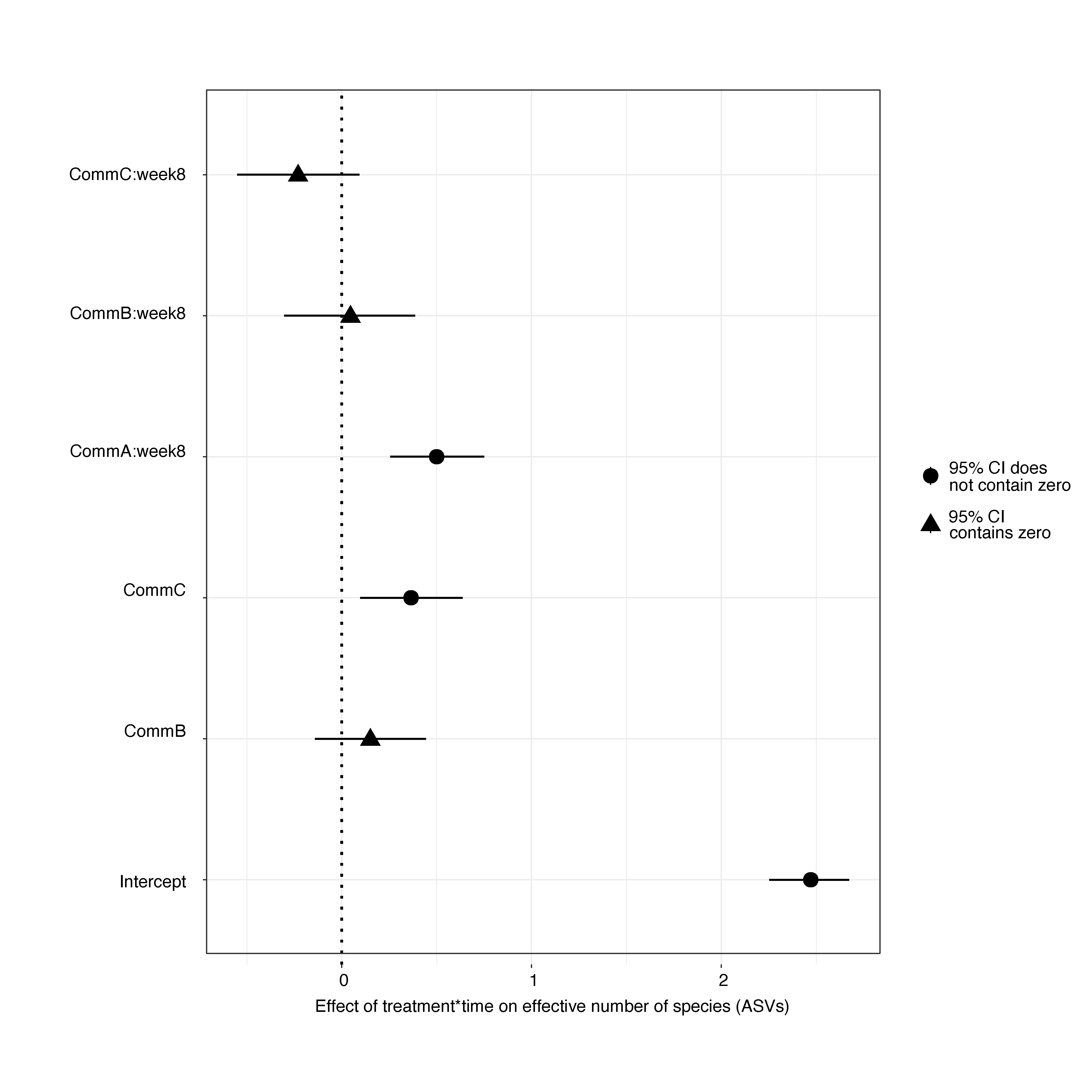

Figure S5. Differences in alpha diversity between bacterial communities. Posterior parameter estimates for the effects of treatment and time on the effective number of species (ASVs). Symbols represent the median parameter estimates and lines represent the 95% credible intervals for the parameter estimate. Parameters with 95% credibility intervals that did not include zero (dashed vertical line) were considered nonzero effects on the response (circles vs. triangles), CommA and week 1 are set as the baseline predictor.

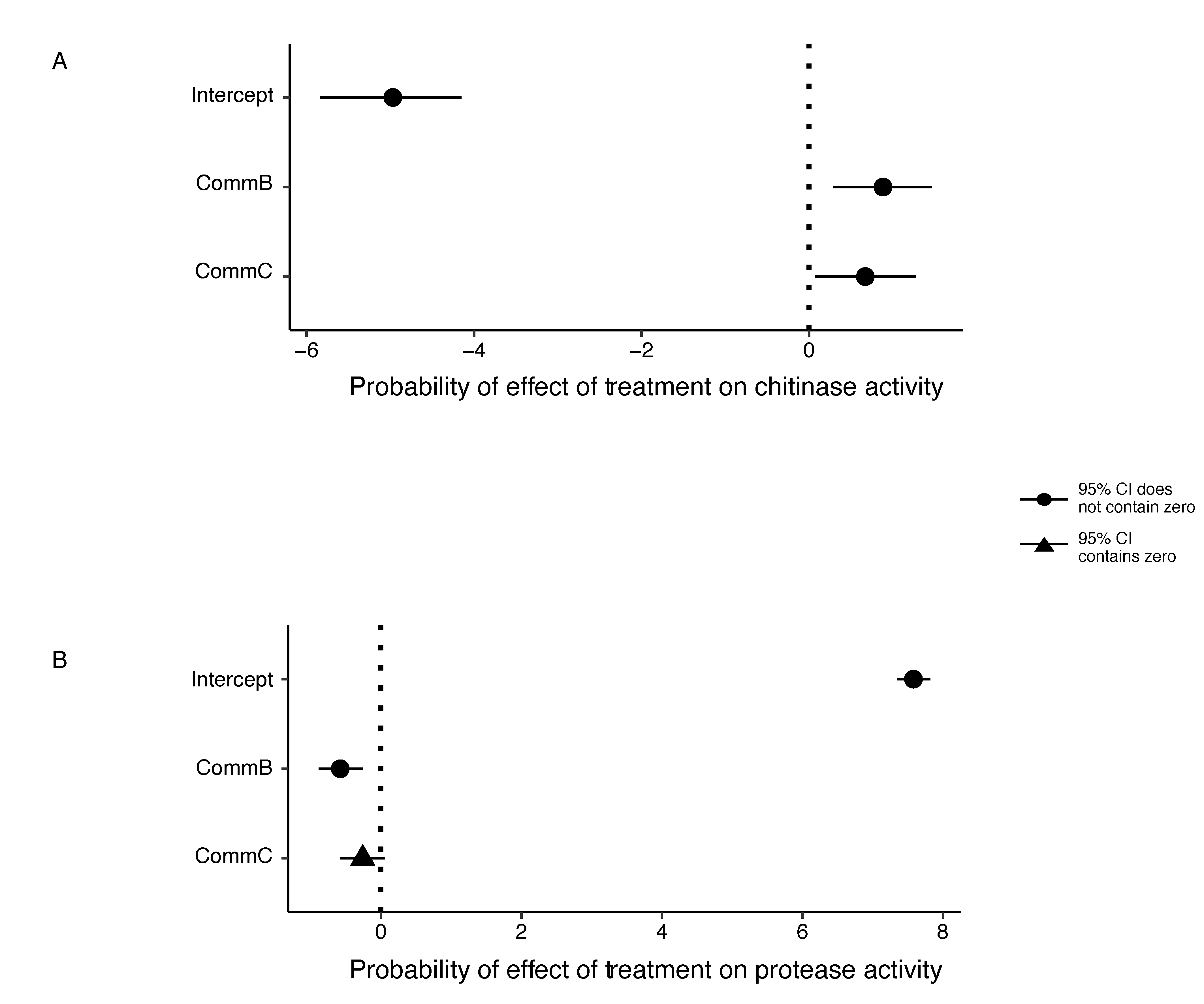

Figure S6. Posterior parameter estimates for the effects of treatment on chitinase and protease activity. Subset to just those plants whose communities were sequenced for metatranscriptomics (3 plants per treatment). Symbols represent the median parameter estimates and lines represent the 95% credible intervals for the parameter estimate. Parameters with 95% credibility intervals that did not include zero (dashed vertical line) were considered nonzero effects on the response (circles vs. triangles). CommA was set as the baseline predictor and time was included in the model as a random intercept. CommB and CommC had higher chitinase activity compared to CommA; CommB had higher protease activity than both CommA and CommC.

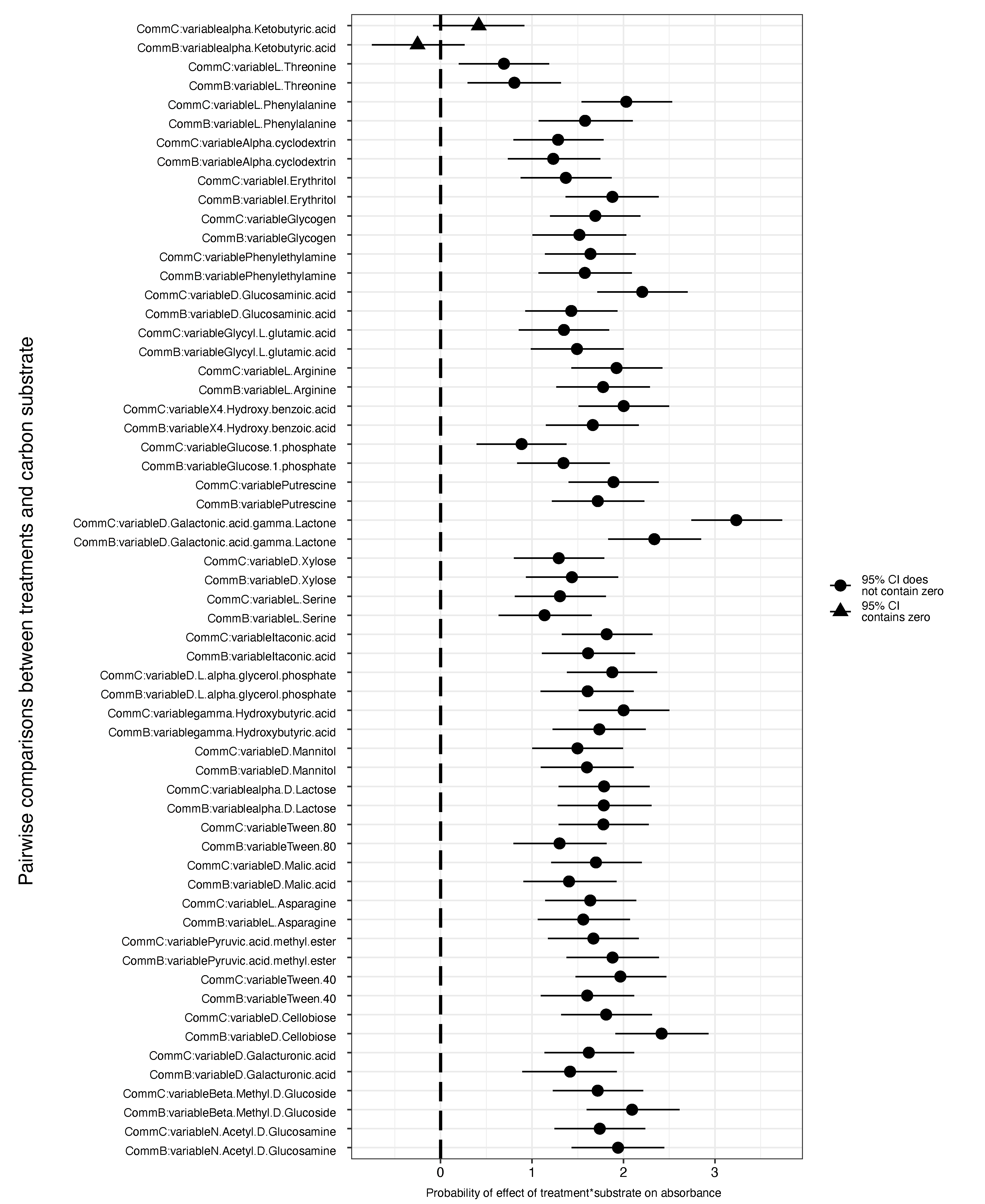

Figure S7. Posterior parameter estimates for the effects of treatment and substrate on absorbance. Symbols represent the median parameter estimates and lines represent the 95% Cis for the parameter estimate. Parameters with 95% credibility intervals that did not include zero (dashed vertical line) were considered nonzero effects on the response (circles vs. triangles). CommA and carbon substrate X2.Hydroxy.benzoic.acid were set as the baseline predictors.

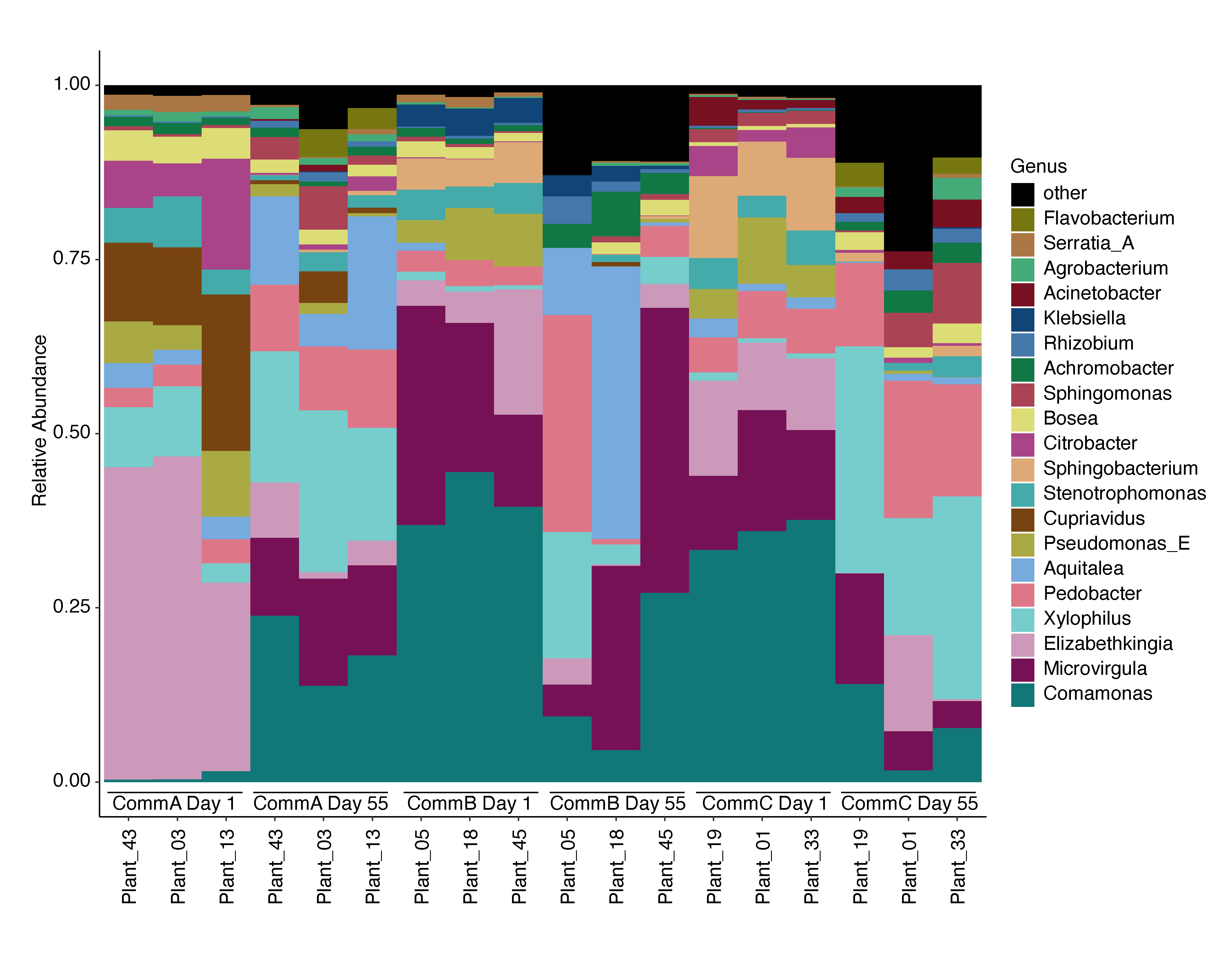

Figure S8. Bacterial relative abundance based on metagenomic analysis. Taxonomic composition differences between the three microbial community treatments identified from shotgun metagenomic analysis of pitcher fluid samples. Relative abundance to the top 20 most abundant genera at day 1 and day 55 for each of the three bacterial community treatments. Taxonomy assigned using the GTDB database based on kmer=31.

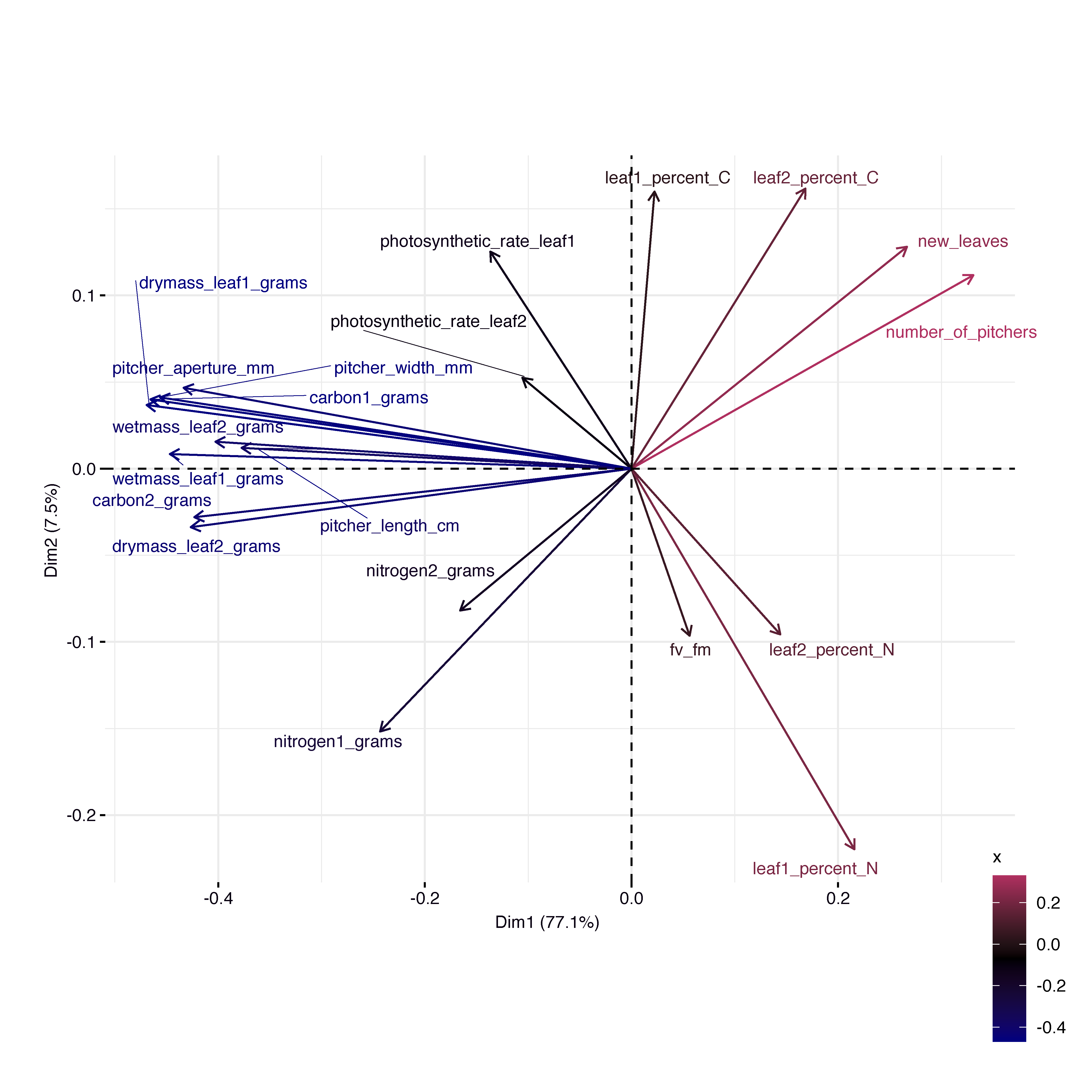

Figure S9. Correlation of plant traits. Principal components analysis (PCA) of scaled plant traits measured for each pitcher (target pitcher = leaf1 and the next youngest pitcher = leaf2 on each plant). The color of the arrows represents the correlation with Dimension (Dim) 1 (x-axis). Dimension 1 explains 77.1% of the variance and dimension 2 explains 7.9% for a total of 85% explained. Along dimension 1, pitcher morphological measures (length, width, aperture, biomass) were tightly correlated with each other so pitcher biomass (grams) was selected to represent plant size and pitcher nitrogen content (mg) represented plant traits that varied along dimension 2 and did not strongly correlate.

**Supplementary Tables**

**Table S1. Pairwise comparisons in Biolog EcoPlate physiological profiles at day 1 and 55 for pitcher bacterial communities (CommA, CommB, CommC), plates read after 72 hours.**

com5 ~ treatment * day, strata = plant, permutations 999

| CommC vs. CommA |  | Df | F | Pr(>F) |
| --- | --- | --- | --- | --- |
|  | Treatment | 1 | 9.9688 | 0.001 |
|  | Day | 1 | 10.0143 | 0.001 |
|  | Treatment:Day | 1 | 4.0190 | 0.014 |
| CommC vs. CommB |  |  |  |  |
|  | Treatment | 1 | 9.8677 | 0.001 |
|  | Day | 1 | 6.7874 | 0.002 |
|  | Treatment:Day | 1 | 2.0407 | 0.090 |
| CommA vs. CommB |  |  |  |  |
|  | Treatment | 1 | 7.8048 | 0.004 |
|  | Day | 1 | 6.7125 | 0.004 |
|  | Treatment:Day | 1 | 3.5927 | 0.023 |

**Table S2. Unweighted UniFrac pairwise comparisons based on 16S amplicon sequencing**.

pairwise.adonis2(uu.dist.16s.treat ~ treatment + day, strata = Null, permutations 999)

| CommC vs. CommA |  | Df | R^2^ | F | Pr(>F) |
| --- | --- | --- | --- | --- | --- |
|  | Treatment | 1 | 0.27025 | 61.2079 | 0.001 |
|  | Day | 7 | 0.08510 | 2.7535 | 0.001 |
| CommC vs. CommB |  |  |  |  |  |
|  | Treatment | 1 | 0.14483 | 26.7313 | 0.001 |
|  | Day | 7 | 0.14542 | 3.8345 | 0.001 |
| CommA vs. CommB |  |  |  |  |  |
|  | Treatment | 1 | 0.26353 | 53.9211 | 0.001 |
|  | Day | 7 | 0.10113 | 2.9559 | 0.001 |

**Table S3. Using analysis of compositions of microbiomes with bias correction (ANCOM-BC), we identified 38 differentially abundant taxa between our treatments and plotted these estimates along with their estimated error.** lfc=log fold change, DA=differentially abundant, se=standard error. Ancombc2(data = df, assay_name = "counts", tax_level = NULL, fix_formula = "treatment", rand_formula = "(1 | day)", p_adj_method = "fdr", prv_cut = 0.3, group="treatment", alpha = 0.05, global = TRUE, pairwise = TRUE)

| ASV | lfc_CommA | lfc_CommC | se_CommA | se_CommC | DA_CommA | DA_CommC |
| --- | --- | --- | --- | --- | --- | --- |
| ASV18 | 2.28088287 | 0 | 0.85764661 | 0.39151026 | TRUE | FALSE |
| ASV2 | 0.85647721 | 0.75524822 | 0.3262171 | 0.33665869 | TRUE | FALSE |
| ASV28 | -0.6789646 | -0.4036594 | 0.23550739 | 0.2442019 | TRUE | FALSE |
| ASV65 | -1.3745555 | 0 | 0.48727129 | 0.39151026 | TRUE | FALSE |
| ASV4 | 0.82878776 | 0.71886 | 0.22909797 | 0.23958411 | TRUE | TRUE |
| ASV9 | 0.07055617 | -0.7617091 | 0.23875579 | 0.25316065 | FALSE | TRUE |
| ASV19 | -2.9715466 | -0.5553903 | 0.79710055 | 0.23307774 | TRUE | FALSE |
| ASV1 | -0.8875793 | -0.1959028 | 0.30478657 | 0.30676597 | TRUE | FALSE |
| ASV11 | 0.4710853 | 3.7877542 | 0.45936395 | 0.432524 | FALSE | TRUE |
| ASV38 | 2.33324976 | 2.64994413 | 0.46725961 | 0.4747444 | TRUE | TRUE |
| ASV34 | 0 | -1.1127227 | 0.38320487 | 0.39420829 | FALSE | TRUE |
| ASV12 | -2.3953791 | -0.1139894 | 0.30102857 | 0.29387924 | TRUE | FALSE |
| ASV8 | -1.1316007 | -0.7806544 | 0.27316502 | 0.28154805 | TRUE | TRUE |
| ASV15 | -0.1816149 | 2.51234793 | 0.31793415 | 0.27634194 | FALSE | TRUE |
| ASV84 | -0.8745872 | -0.0362883 | 0.27831542 | 0.25077011 | TRUE | FALSE |
| ASV57 | -0.9444645 | -0.0494317 | 0.2862049 | 0.24735335 | TRUE | FALSE |
| ASV106 | -0.8664169 | -0.078243 | 0.26386658 | 0.22958842 | TRUE | FALSE |
| ASV67 | 1.26668178 | 2.33520807 | 0.32291121 | 0.31662338 | TRUE | TRUE |
| ASV21 | 0.20373376 | -0.8973839 | 0.28504718 | 0.29655565 | FALSE | TRUE |
| ASV59 | -0.7356318 | -1.0637647 | 0.22144103 | 0.23245335 | TRUE | TRUE |
| ASV5 | 1.01782947 | -0.5900468 | 0.25855364 | 0.26786661 | TRUE | FALSE |
| ASV6 | -1.0924291 | -0.7822658 | 0.23655704 | 0.23814965 | TRUE | TRUE |
| ASV31 | -1.967328 | -1.3175785 | 0.32111856 | 0.27845587 | TRUE | TRUE |
| ASV30 | 0.02063799 | -1.1920085 | 0.26247751 | 0.27151989 | FALSE | TRUE |
| ASV88 | -1.2316125 | -1.1584666 | 0.37293824 | 0.31068293 | TRUE | TRUE |
| ASV10 | 2.48587669 | 3.87683718 | 0.77798985 | 0.70223672 | TRUE | TRUE |
| ASV121 | -1.5892854 | -0.1279019 | 0.30353483 | 0.23251559 | TRUE | FALSE |
| ASV69 | -0.7396856 | -1.5488697 | 0.20033138 | 0.21402957 | TRUE | TRUE |
| ASV90 | -1.0598345 | -0.3681058 | 0.24679575 | 0.25811064 | TRUE | FALSE |
| ASV123 | -0.895807 | -0.1304857 | 0.36140543 | 0.27778054 | TRUE | FALSE |
| ASV85 | -2.5631267 | -0.697296 | 0.38623488 | 0.28440992 | TRUE | FALSE |
| ASV32 | -2.6922573 | 0.32432347 | 0.32897822 | 0.23346745 | TRUE | FALSE |
| ASV41 | -1.7207563 | 0.77619731 | 0.72608583 | 0.30248531 | TRUE | TRUE |
| ASV62 | -2.1179012 | -0.3840785 | 0.42798943 | 0.26251152 | TRUE | FALSE |
| ASV35 | -1.3413746 | 0.42010938 | 0.30682567 | 0.27697165 | TRUE | FALSE |
| ASV58 | -1.6998289 | 0.8386147 | 0.42003553 | 0.27163856 | TRUE | TRUE |
| ASV46 | -1.0877948 | 0.00138532 | 0.35357518 | 0.34144047 | TRUE | FALSE |
| ASV39 | -0.6185634 | 0.25422089 | 0.2425294 | 0.3216076 | TRUE | FALSE |

**Table S4. MAG taxonomy and quality metrics**.

| **Name** | **new_bin_name** | **Completeness** | **Contamination** | **quality_grade** | **Contig_N50** | **Average_Gene_Length** | **Genome_Size** | **GC_Content** | **Total_Coding_Sequences** | **lineage** | **query_md5** | **f_weighted_at_rank** | **bp_match_at_rank** | **query_ani_at_rank** |
| --- | --- | --- | --- | --- | --- | --- | --- | --- | --- | --- | --- | --- | --- | --- |
| maxbin.41 | MAG_34 | 99.99 | 10.05 | discard | 117302 | 304.1812541 | 6320792 | 0.62 | 6124 | d__Bacteria; p__Pseudomonadota; c__Gammaproteobacteria;o__Burkholderiales;f__Burkholderiaceae_B;g__Comamonas;s__Comamonas testosteroni_C | 2503fe2d | 0.245283019 | 1560000 | 0.897695969 |
| concoct.81_sub | MAG_27 | 90.82 | 10.74 | discard | 10693 | 276.216608 | 6750378 | 0.6 | 7105 | d__Bacteria; p__Pseudomonadota; c__Alphaproteobacteria;o__Rhizobiales;f__Rhizobiaceae;g__Agrobacterium;s__Agrobacterium fabacearum | 7ea57366 | 0.10521701 | 720000 | 0.996175442 |
| concoct.74 | MAG_44 | 86.49 | 11.21 | discard | 4326 | 256.3423595 | 5575643 | 0.54 | 6315 | d__Bacteria; p__Pseudomonadota; c__Gammaproteobacteria;o__Enterobacterales;f__Enterobacteriaceae;g__Serratia_A;s__Serratia_A fonticola | 62bbd086 | 0.943229836 | 5134000 | 0.871970498 |
| concoct.70_sub | MAG_11 | 73.94 | 21.93 | discard | 2301 | 228.6382245 | 3372305 | 0.46 | 4348 | d__Bacteria; p__Bacillota_C; c__Negativicutes;o__Sporomusales_C;f__DSM-15969;g__Anaerospora;s__Anaerospora sp002337925 | 900253b4 | 0.032017408 | 103000 | 0.93483343 |
| metabat2.44_sub | MAG_47 | 88.45 | 35.8 | discard | 5952 | 301.0039273 | 6906112 | 0.68 | 6875 | d__Bacteria; p__Pseudomonadota; c__Gammaproteobacteria;o__Xanthomonadales;f__Rhodanobacteraceae;g__Dokdonella_A;s__Dokdonella_A sp017744955 | 2a7724f1 | 0.014307229 | 95000 | 0.859142103 |
| concoct.3 | MAG_19 | 100 | 37.58 | discard | 137612 | 339.2890728 | 8413010 | 0.41 | 7614 | d__Bacteria; p__Bacteroidota; c__Bacteroidia;o__Sphingobacteriales;f__Sphingobacteriaceae;g__Pedobacter;s__Pedobacter nutrimenti | 6b7d278f | 0.409965348 | 3431000 | 0.822039738 |
| concoct.84 | MAG_48 | 99.82 | 73.45 | discard | 40959 | 328.1228387 | 8471315 | 0.67 | 7750 | d__Bacteria; p__Pseudomonadota; c__Gammaproteobacteria;o__Xanthomonadales;f__Xanthomonadaceae;g__Stenotrophomonas;s__Stenotrophomonas sp002192255 | 26e87d98 | 0.088797302 | 711000 | 0.829564005 |
| metabat2.12 | MAG_45 | 100 | 0.04 | high | 269072 | 312.4638309 | 3919568 | 0.39 | 3691 | d__Bacteria; p__Pseudomonadota; c__Gammaproteobacteria;o__Pseudomonadales;f__Moraxellaceae;g__Acinetobacter;s__Acinetobacter seifertii | 12354c55 | 0.88215998 | 3496000 | 0.953139318 |
| maxbin.11 | MAG_01 | 100 | 0.06 | high | 74848 | 323.2122769 | 3887567 | 0.71 | 3698 | d__Bacteria; p__Actinomycetota; c__Actinomycetia;o__Actinomycetales;f__Microbacteriaceae;g__Leifsonia;s__Leifsonia sp001898805 | 5a696026 | 0.116290984 | 454000 | 0.901116999 |
| concoct.92_sub | MAG_16 | 95.5 | 0.13 | high | 42866 | 333.9214864 | 3678956 | 0.37 | 3337 | d__Bacteria; p__Bacteroidota; c__Bacteroidia;o__Flavobacteriales;f__Flavobacteriaceae;g__Flavobacterium;s__Flavobacterium microcysteis | 1f669bfb | 0.128183832 | 463000 | 0.890346875 |
| metabat2.91 | MAG_02 | 99.77 | 0.16 | high | 336160 | 326.6919406 | 1988347 | 0.63 | 1886 | d__Bacteria; p__Actinomycetota; c__Actinomycetia;o__Actinomycetales;f__Microbacteriaceae;g__Leifsonia;s__Leifsonia sp009765225 | e99fa88c | 0.00305033 | 6000 | 0.989066875 |
| maxbin.10 | MAG_09 | 98.42 | 0.35 | high | 45578 | 304.6598456 | 2732790 | 0.46 | 2590 | d__Bacteria; p__Bacillota_A; c__Clostridia;o__Oscillospirales;f__Acutalibacteraceae;g__Caproiciproducens;s__Caproiciproducens sp002338255 | a858386e | 0.00795756 | 21000 | 0.875708865 |
| concoct.7 | MAG_39 | 100 | 0.53 | high | 37177 | 317.9608681 | 6190281 | 0.67 | 5852 | d__Bacteria; p__Pseudomonadota; c__Gammaproteobacteria;o__Burkholderiales;f__Burkholderiaceae_C;g__Achromobacter;s__Achromobacter sp016428805 | 41529c77 | 0.877144707 | 5419000 | 0.971645788 |
| concoct.20 | MAG_22 | 90.9 | 0.64 | high | 69468 | 341.2928881 | 4401942 | 0.44 | 3923 | d__Bacteria; p__Bacteroidota; c__Bacteroidia;o__Sphingobacteriales;f__Sphingobacteriaceae;g__Pedobacter;s__Pedobacter sp016429155 | c2f378b7 | 0.711205929 | 3167000 | 0.846410758 |
| metabat2.80 | MAG_14 | 98.85 | 0.68 | high | 149806 | 341.1593153 | 5297186 | 0.46 | 4557 | d__Bacteria; p__Bacteroidota; c__Bacteroidia;o__Chitinophagales;f__Chitinophagaceae;g__Edaphocola;s__Edaphocola koreensis | 81013bce | 0.002299732 | 12000 | 0.99840909 |
| concoct.41 | MAG_17 | 100 | 0.99 | high | 381815 | 322.5015819 | 4452723 | 0.36 | 4109 | d__Bacteria; p__Bacteroidota;c__Bacteroidia;o__Flavobacteriales;f__Weeksellaceae;g__Elizabethkingia;s__Elizabethkingia miricola | a60dd1d3 | 0.975498054 | 4260000 | 0.996899806 |
| concoct.48_sub | MAG_23 | 100 | 1.05 | high | 640265 | 343.4683262 | 6769677 | 0.39 | 5825 | d__Bacteria; p__Bacteroidota;c__Bacteroidia;o__Sphingobacteriales;f__Sphingobacteriaceae;g__Sphingobacterium;s__Sphingobacterium siyangense | 8ae0385f | 0.449198647 | 3055000 | 0.999200091 |
| concoct.60 | MAG_31 | 92.66 | 1.22 | high | 22989 | 306.3097527 | 4429949 | 0.68 | 4368 | d__Bacteria; p__Pseudomonadota;c__Alphaproteobacteria;o__Sphingomonadales;f__Sphingomonadaceae;g__Sphingomonas;s__Sphingomonas sp017418975 | 95e86b8b | 0.712693358 | 3133000 | 0.857637285 |
| concoct.83 | MAG_26 | 92.42 | 1.22 | high | 174707 | 315.8816997 | 4397229 | 0.59 | 4142 | d__Bacteria; p__Pseudomonadota;c__Alphaproteobacteria;o__Rhizobiales;f__Rhizobiaceae;g__Agrobacterium;s__Agrobacterium fabacearum | 9e7e1912 | 0.900923215 | 4001000 | 0.97451457 |
| concoct.82_sub | MAG_21 | 99.88 | 1.3 | high | 672634 | 349.3753887 | 5332767 | 0.39 | 4502 | d__Bacteria; p__Bacteroidota;c__Bacteroidia;o__Sphingobacteriales;f__Sphingobacteriaceae;g__Pedobacter;s__Pedobacter sp016429065 | 65ea6890 | 0.978470676 | 5272000 | 0.947240999 |
| maxbin.12_sub | MAG_10 | 100 | 1.63 | high | 332405 | 308.1145038 | 4483247 | 0.42 | 4192 | d__Bacteria; p__Bacillota_C;c__Negativicutes;o__Propionisporales;f__DSM-13327;g__Pelosinus;s__Pelosinus fermentans_A | be376984 | 0.005874379 | 26000 | 0.992992627 |
| metabat2.65 | MAG_04 | 99.82 | 1.64 | high | 142208 | 324.0040777 | 4514432 | 0.64 | 4169 | d__Bacteria; p__Actinomycetota;c__Actinomycetia;o__Mycobacteriales;f__Mycobacteriaceae;g__Gordonia;s__Gordonia polyisoprenivorans | 5ee9f234 | 0.009036808 | 41000 | 0.991881831 |
| concoct.78 | MAG_24 | 98.96 | 1.99 | high | 15182 | 289.8501554 | 4635018 | 0.68 | 4825 | d__Bacteria; p__Pseudomonadota;c__Alphaproteobacteria;o__Caulobacterales;f__Caulobacteraceae;g__Phenylobacterium;s__Phenylobacterium sp013822795 | e62569ec | 0.059270517 | 273000 | 0.894925021 |
| maxbin.54 | MAG_18 | 99.89 | 2.02 | high | 208541 | 326.176505 | 3690367 | 0.34 | 3422 | d__Bacteria; p__Bacteroidota;c__Bacteroidia;o__Flavobacteriales;f__Weeksellaceae;g__Epilithonimonas;s__Epilithonimonas hispanica | ee598531 | 0.02406344 | 88000 | 0.998116444 |
| concoct.77_sub | MAG_15 | 95.43 | 2.03 | high | 19859 | 327.5583248 | 5424429 | 0.34 | 4895 | d__Bacteria; p__Bacteroidota;c__Bacteroidia;o__Flavobacteriales;f__Flavobacteriaceae;g__Flavobacterium;s__Flavobacterium chilense | b41e366b | 0.968813811 | 5219000 | 0.998978495 |
| concoct.35 | MAG_43 | 100 | 2.04 | high | 173044 | 314.8735437 | 5020425 | 0.52 | 4721 | d__Bacteria; p__Pseudomonadota;c__Gammaproteobacteria;o__Enterobacterales;f__Enterobacteriaceae;g__Citrobacter;s__Citrobacter braakii | 0f1b4e11 | 0.951840796 | 4783000 | 0.912880966 |
| maxbin.21 | MAG_36 | 92.78 | 2.81 | high | 274989 | 317.8978723 | 3584135 | 0.47 | 3525 | d__Bacteria; p__Pseudomonadota;c__Gammaproteobacteria;o__Burkholderiales;f__Burkholderiaceae_B;g__Variovorax;s__Variovorax sp003019815 | 0e6d1763 | 0.001114827 | 4000 | 0.929938837 |
| concoct.58 | MAG_25 | 93.07 | 3.21 | high | 8686 | 279.935212 | 3619376 | 0.69 | 3797 | d__Bacteria; p__Pseudomonadota;c__Alphaproteobacteria;o__Rhizobiales;f__Ancalomicrobiaceae;g__Pinisolibacter;s__Pinisolibacter sp002298965 | aef23108 | 0.186326702 | 665000 | 0.999298167 |
| concoct.95 | MAG_46 | 100 | 3.44 | high | 27401 | 310.4301915 | 7101217 | 0.63 | 6790 | d__Bacteria; p__Pseudomonadota;c__Gammaproteobacteria;o__Pseudomonadales;f__Pseudomonadaceae;g__Pseudomonas_E;s__Pseudomonas_E fluorescens_AP | f14046b1 | 0.814279703 | 5805000 | 0.996640004 |
| concoct.69 | MAG_42 | 99.93 | 3.45 | high | 99600 | 321.2930811 | 4189321 | 0.61 | 3859 | d__Bacteria; p__Pseudomonadota;c__Gammaproteobacteria;o__Burkholderiales;f__Chromobacteriaceae;g__Aquitalea;s__Aquitalea magnusonii | 5a4d1c90 | 0.776708373 | 3228000 | 0.924863014 |
| metabat2.16_sub | MAG_29 | 92.53 | 3.82 | high | 150427 | 313.0756176 | 4860522 | 0.62 | 4655 | d__Bacteria; p__Pseudomonadota;c__Alphaproteobacteria;o__Rhizobiales;f__Rhizobiaceae;g__Shinella;s__Shinella sumterensis | 9efb1a37 | 0.035237698 | 169000 | 0.99339445 |
| concoct.38 | MAG_38 | 97.02 | 4.1 | high | 28539 | 313.8317801 | 4416768 | 0.69 | 4292 | d__Bacteria; p__Pseudomonadota;c__Gammaproteobacteria;o__Burkholderiales;f__Burkholderiaceae_B;g__Xylophilus;s__Xylophilus sp016428875 | 3e768fb3 | 0.908232119 | 4038000 | 0.932945607 |
| concoct.113 | MAG_28 | 100 | 4.12 | high | 227871 | 337.2025518 | 4202342 | 0.61 | 3762 | d__Bacteria; p__Pseudomonadota;c__Alphaproteobacteria;o__Rhizobiales;f__Rhizobiaceae;g__Agrobacterium;s__Agrobacterium fabacearum | 8c56278f | 0.027310924 | 117000 | 0.847287951 |
| concoct.46 | MAG_20 | 66.93 | 0.94 | medium | 4181 | 268.6341524 | 3668623 | 0.43 | 4029 | d__Bacteria; p__Bacteroidota;c__Bacteroidia;o__Sphingobacteriales;f__Sphingobacteriaceae;g__Pedobacter;s__Pedobacter sp016429005 | 30d07701 | 0.008558807 | 31000 | 0.838541929 |
| concoct.30_sub | MAG_05 | 52.85 | 1.26 | medium | 1868 | 232.3076923 | 2450838 | 0.62 | 3159 | d__Bacteria; p__Actinomycetota;c__Actinomycetia;o__Propionibacteriales;f__Nocardioidaceae;g__Nocardioides;s__Nocardioides sp018831735 | 856a4314 | 0.005688744 | 14000 | 0.988546654 |
| concoct.25 | MAG_13 | 81.72 | 1.29 | medium | 4866 | 305.9399027 | 6460613 | 0.46 | 6373 | d__Bacteria; p__Bacteroidota;c__Bacteroidia;o__Chitinophagales;f__Chitinophagaceae;g__Chitinophaga;s__Chitinophaga hostae | e2eb9d22 | 0.016336241 | 103000 | 0.830741786 |
| concoct.80 | MAG_37 | 80.05 | 1.38 | medium | 7181 | 269.4011252 | 4247672 | 0.69 | 4799 | d__Bacteria; p__Pseudomonadota;c__Gammaproteobacteria;o__Burkholderiales;f__Burkholderiaceae_B;g__Variovorax;s__Variovorax sp009765735 | 0226b0c4 | 0.025659647 | 106000 | 0.989133535 |
| metabat2.47 | MAG_12 | 87.12 | 1.59 | medium | 37471 | 353.0781985 | 4686069 | 0.44 | 4041 | d__Bacteria; p__Bacteroidota;c__Bacteroidia;o__Bacteroidales;f__Bacteroidaceae;g__Bacteroides;s__Bacteroides neonati | d0d1332a | 0.123811582 | 573000 | 0.992547534 |
| concoct.67_sub | MAG_06 | 51.55 | 1.65 | medium | 1729 | 215.8222381 | 2885276 | 0.72 | 4191 | d__Bacteria; p__Actinomycetota;c__Actinomycetia;o__Propionibacteriales;f__Nocardioidaceae;g__Nocardioides;s__Nocardioides sp018831735 | 821b2eea | 0.79303208 | 2299000 | 0.995780434 |
| concoct.101_sub | MAG_08 | 64.12 | 2.63 | medium | 3769 | 266.5700991 | 3034594 | 0.41 | 3331 | d__Bacteria; p__Bacillota_A;c__Clostridia;o__Lachnospirales;f__Lachnospiraceae;g__Anaerocolumna;s__Anaerocolumna xylanovorans | 43bc704f | 0.039647577 | 117000 | 0.888557436 |
| metabat2.4_sub | MAG_33 | 86.84 | 3.75 | medium | 19306 | 323.5213102 | 5406569 | 0.67 | 5068 | d__Bacteria; p__Pseudomonadota;c__Gammaproteobacteria;o__Burkholderiales;f__Burkholderiaceae_B;g__Comamonas;s__Comamonas acidovorans | c2a411d8 | 0.887995512 | 4749000 | 0.935880562 |
| concoct.37_sub | MAG_41 | 80.77 | 3.87 | medium | 4003 | 251.9963186 | 2535214 | 0.56 | 2988 | d__Bacteria; p__Pseudomonadota;c__Gammaproteobacteria;o__Burkholderiales;f__Chromobacteriaceae;g__Aquitalea;s__Aquitalea magnusonii | f25d729c | 0.004258614 | 11000 | 0.855624522 |
| concoct.68 | MAG_32 | 100 | 5.48 | medium | 113119 | 327.5562357 | 4266263 | 0.64 | 3921 | d__Bacteria; p__Pseudomonadota;c__Gammaproteobacteria;o__Burkholderiales;f__Aquaspirillaceae;g__Microvirgula;s__Microvirgula aerodenitrificans | 5cc27572 | 0.804130889 | 3465000 | 0.960872991 |
| concoct.57 | MAG_07 | 78.52 | 6.44 | medium | 2956 | 230.6606796 | 3432409 | 0.31 | 4120 | d__Bacteria; p__Bacillota_A;c__Clostridia;o__Clostridiales;f__Clostridiaceae;g__Clostridium_J;s__Clostridium_J tunisiense | 715ce2fa | 0.003187482 | 11000 | 0.846547946 |
| concoct.56 | MAG_40 | 99.99 | 6.68 | medium | 41167 | 311.8115553 | 9589839 | 0.65 | 8931 | d__Bacteria; p__Pseudomonadota;c__Gammaproteobacteria;o__Burkholderiales;f__Burkholderiaceae;g__Cupriavidus;s__Cupriavidus basilensis | 5112c95a | 0.699700663 | 6545000 | 0.803061177 |
| maxbin.17 | MAG_03 | 98.98 | 8.55 | medium | 17934 | 291.7331334 | 3844954 | 0.69 | 4002 | d__Bacteria; p__Actinomycetota;c__Actinomycetia;o__Actinomycetales;f__Microbacteriaceae;g__Microbacterium;s__Microbacterium algeriense | a5db894d | 0.005717397 | 21000 | 0.955678585 |
| maxbin.15_sub | MAG_30 | 95.01 | 9.47 | medium | 48074 | 320.1657938 | 5832709 | 0.66 | 5537 | d__Bacteria; p__Pseudomonadota;c__Alphaproteobacteria;o__Sphingomonadales;f__Sphingomonadaceae;g__Sphingomonas;s__Sphingomonas sp017304125 | d2b5a109 | 0.290165361 | 1667000 | 0.886718425 |
| concoct.1_sub | MAG_35 | 67.99 | 9.68 | medium | 1695 | 213.4947754 | 5107203 | 0.69 | 7369 | d__Bacteria; p__Pseudomonadota;c__Gammaproteobacteria;o__Burkholderiales;f__Burkholderiaceae_B;g__Variovorax;s__Variovorax sp003019815 | dfc029c7 | 0.225864481 | 1130000 | 0.995963592 |
